# Aggregation state of *Mycobacterium tuberculosis* impacts host immunity and augments pulmonary disease pathology

**DOI:** 10.1101/2021.05.19.444830

**Authors:** Afsal Kolloli, Ranjeet Kumar, Pooja Singh, Anshika Narang, Gilla Kaplan, Alex Sigal, Selvakumar Subbian

**Author notes:** **Correspondence**: Selvakumar Subbian, PHRI/ICPH Center, 225 Warren Street, Room W310.W, Newark, NJ07103, USA.

## Abstract

Phagocytosis of *Mycobacterium tuberculosis* (Mtb) aggregates, rather than similar numbers of single bacilli, induces host macrophage death and favors bacterial growth. Here, we examined whether aggregation contributes to enhanced Mtb pathogenicity in vivo in rabbit lungs. Rabbits were exposed to infectious aerosols containing mainly Mtb-aggregates (Mtb-AG) or Mtb-single cells (Mtb-SC). The lung bacterial load, histology, and immune cell composition were investigated over time. Genome-wide transcriptome analysis, cellular and tissue-level assays, and immunofluorescent imaging were performed on lung tissue to define and compare differential immune activation and pathogenesis between Mtb-AG and Mtb-SC infection.

Lung bacillary loads, disease scores, lesion size, and structure were significantly higher in Mtb-AG than in Mtb-SC infected animals. A differential immune cell distribution and activation were noted in the lungs and spleen of the two groups of infected animals. Mtb-AG infected animals also showed early induction of inflammatory network genes associated with necrosis and reduced host cell viability. Consistently, larger lung granulomas with clumped Mtb, extensive necrotic foci, and elevated matrix metalloproteases expression were observed in Mtb-AG infected rabbits. Our findings suggest that bacillary aggregation increases Mtb fitness for improved growth and accelerated lung inflammation and cell death, thereby exacerbating disease pathology in the lungs.

## INTRODUCTION

*Mycobacterium tuberculosis* (Mtb), the causative agent of tuberculosis (TB) has infected about a fourth of the global population (1). Of those exposed to Mtb, about 95% manifest immune-controlled latent infection while the remaining 5% progress to active TB disease (1). The host and pathogen determinants that favor the progression of infection to active disease are complex and not fully understood. Immune suppression, as seen in HIV-infected individuals, is associated with loss of protective immunity and development of full-blown TB disease (2). In addition, exposure to a high dose of viable Mtb, exhaled or coughed by a patient with cavitary pulmonary disease, favors a more efficient spread of the infection and progression to active disease (3,4). Moreover, specific properties of different clinical Mtb strain can also impact their ability to establish infection and progress to disease, as seen both in animal models and clinical studies (4–7).

Mtb has a natural tendency to form aggregates (AG) or clumps (a.k.a “cording”) in broth cultures due to the presence of a waxy cell wall. Electron microscopy studies of pathogenic mycobacteria have shown cording in the lag, log, and stationary growth phases (8, 9). We recently reported that exposure of human monocyte-derived macrophages to clumps/aggregates of Mtb, compared to a similar number of single bacilli, interferes with the control of intracellular bacillary growth and results in efficient replication of the bacilli, in association with the death of the phagocyte (10). Furthermore, efferocytosis of Mtb-loaded-dead macrophages resulted in more host cell death and increased bacterial proliferation (10). It is important to note that Mtb clumps are often seen at the cavitary surface of pulmonary lesions and in the infectious bioaerosols exhaled by patients with active pulmonary TB (9, 11, 12). Based on these observations, we hypothesize that host exposure to Mtb-AG, as opposed to single bacilli (Mtb-SC), may be an effective bacterial strategy for establishing an infection that is more likely to progress to active disease.

To test this hypothesis, we evaluated in vivo infection of rabbits, shown previously to develop TB disease pathogenesis, similar to that seen in humans. Animals were exposed to aerosolized Mtb-AG, and the outcome of infection was compared with animals infected with similar numbers of Mtb-SC. The kinetics of Mtb growth, disease pathology, and immune cell activation were monitored over time to determine whether infection by Mtb-AG results in increased bacillary load and progression to more severe disease in the lungs.

## RESULTS

### Infection with Mtb-AG results in consistently increased Mtb burden in rabbit lungs

To determine the relative growth kinetics of Mtb-AG versus Mtb-SC in vivo, rabbits were infected with aerosolized bacilli, and the lung bacillary load was evaluated over time. At T=0 (3 hrs. post-infection), the number of CFU was similar in both Mtb-AG and Mtb-SC infected rabbit lungs (Figure 1A). A significant increase in the bacillary load was noted in Mtb-AG infected lungs by one week post-infection, compared to Mtb-SC. The differential growth between Mtb-AG and Mtb-SC was sustained at two and four weeks post-infection (Figure 1A). The CFU data was consistent with and supported by the data from imaging fluorescent Mtb and CEQ that measured the net bacillary load in the lungs (Figure 1B and C). Together, these findings suggest that Mtb-AG grows more efficiently than Mtb-SC in vivo in the lungs of infected rabbits.

**Figure 1.**
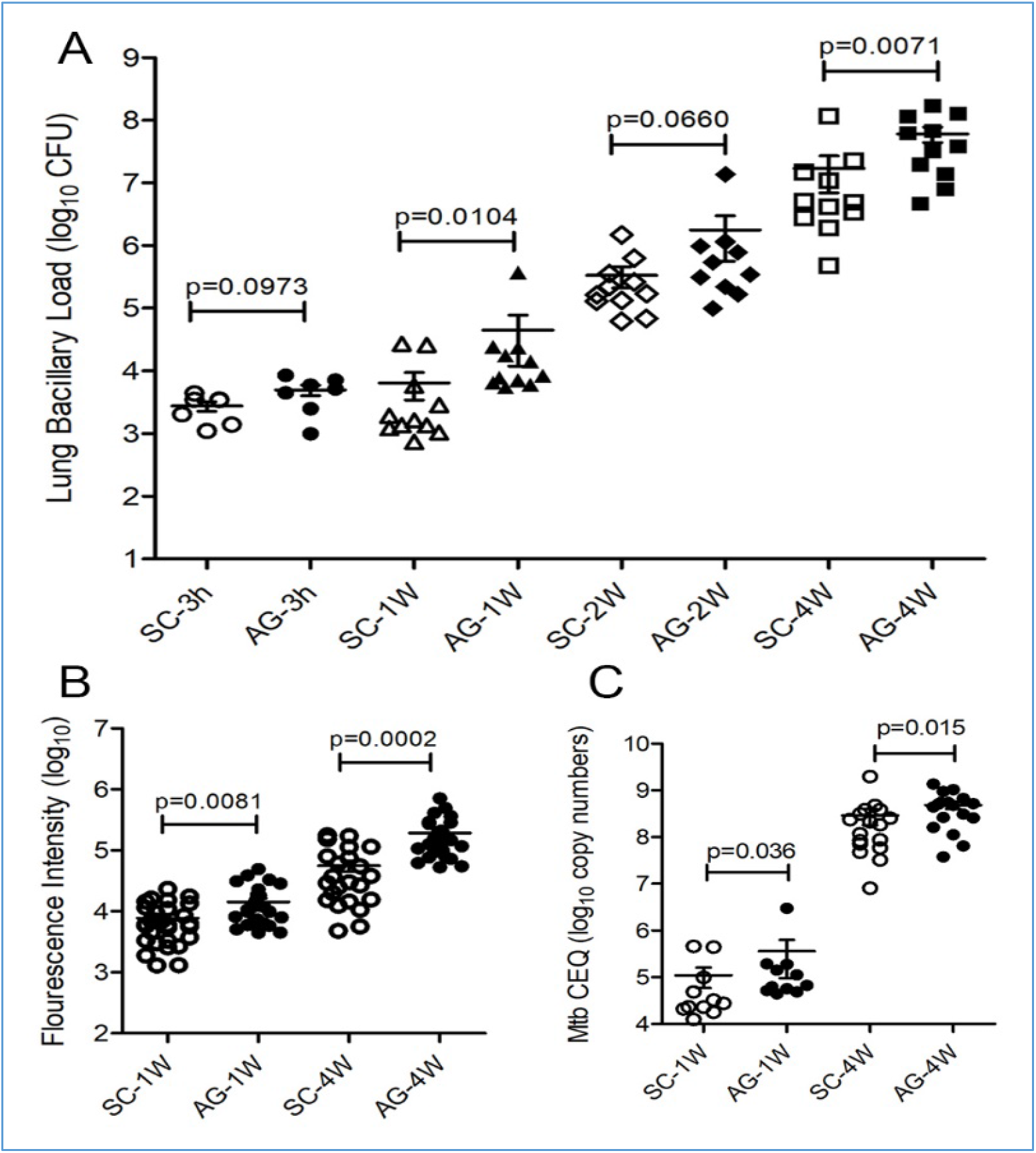
Growth kinetics of Mtb-AG and Mtb-SC in rabbit lungs. **A**. Bacillary load measured as the number of CFU in rabbit lungs infected with Mtb-AG compared to Mtb-SC at 3hrs (T=0), 1, 2 and 4 weeks post-infection. Experiments were performed with n=6-11 per group per time point. p-value was calculated using one-way ANOVA with Tukey’s post-correction for multiple group comparison. **B.** Net bacterial load in the lung sections as quantified by immunofluorescence of Mtb anti-LAM antibody at 1- and 4-week post-infection. Data was compiled from measuring fluorescent intensity in 24-30 fields per lung (n=4 per group per time point). p-value was calculated using Mann-Whitney U-Test. **C.** Total lung bacterial load estimated by Mtb-CEQ analysis at 4 weeks post-infection. Copy number in lung samples were determined from standard curve developed with a known concentration of Mtb genomic DNA isolated from broth cultures. Data compiled from n=3 per group per time point and repeated twice. p-value was calculated using Mann-Whitney U-Test.

### Infection with Mtb-AG exacerbates disease pathology and affects immune cell distribution in the lungs

We next evaluated the granulomatous response in the lungs of Mtb-AG or Mtb-SC infected rabbits. Rabbit lung gross pathology at four weeks post-infection showed larger sub-pleural lesions in the Mtb-AG infected group (Figure 2A and B). Histologic examination of H&E-stained lung sections showed multiple, coalescent lesions with several prominent necrotic foci with abundant polymorphonuclear cell accumulation, preferentially in the Mtb-AG-infected animals; in these lesions, Mtb bacilli were seen mostly as larger aggregates (Figure 2C and E). In contrast, in the Mtb-SC infected lungs, the lesions were less necrotic, and the bacilli were seen as singles or small clumps (Figure 2D and F). Although no significant difference was noted in the number of subpleural granulomas between Mtb-AG and Mtb-SC infected rabbit lungs, the Mtb-AG infected lungs had significantly larger lesions and a higher pulmonary disease score than Mtb-SC infected animals (Figure 2 G-I). These observations suggest that Mtb-AG infection exacerbates pulmonary disease pathology leading to more extensive cell recruitment, inflammation and necrosis than Mtb-SC infection.

**Figure 2.**
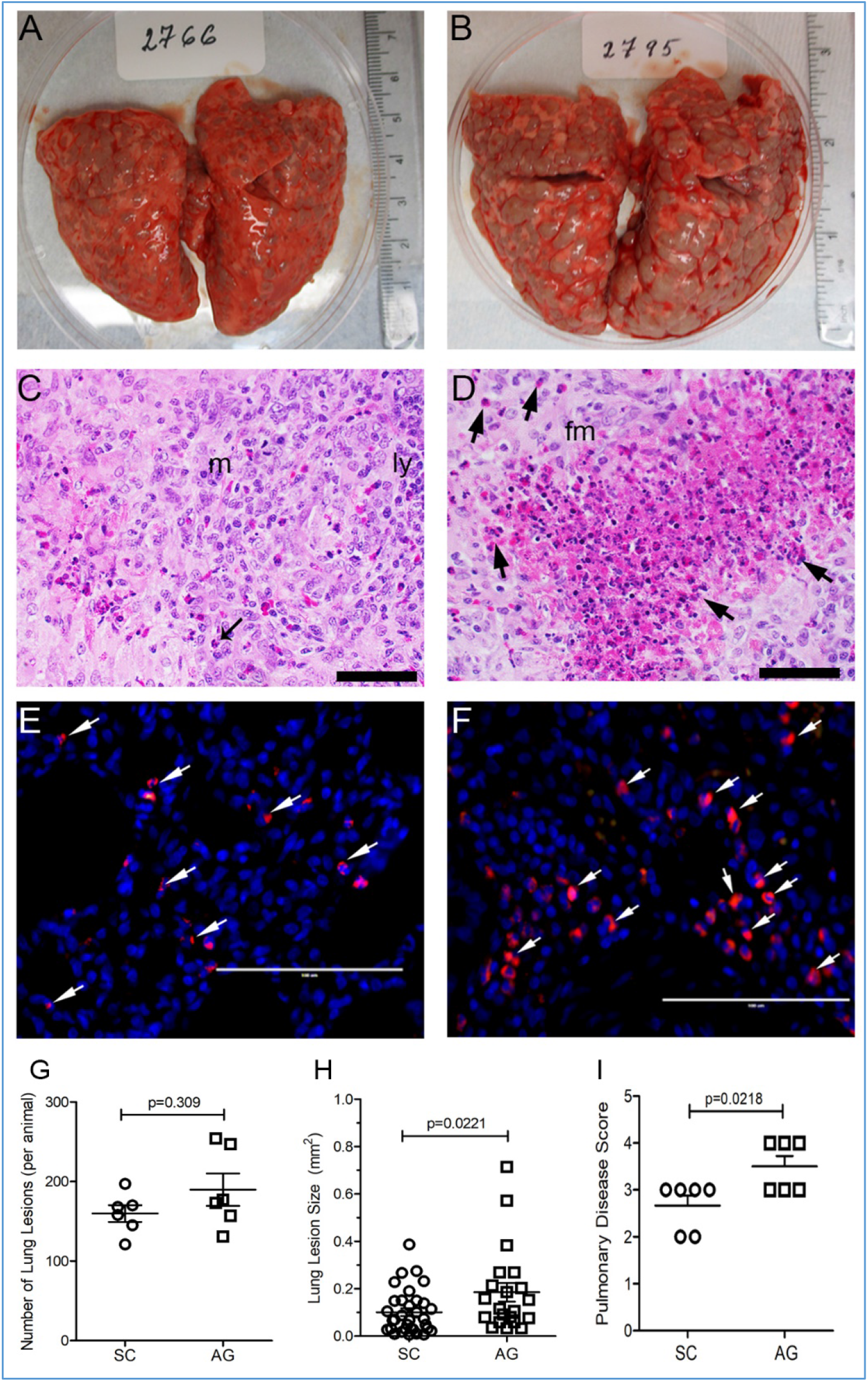
Gross and histopathologic analysis of Mtb-AG or Mtb-SC-infected rabbit lungs at 4 weeks post-infection. A. Gross pathology of Mtb-SC infected rabbit lungs showing multiple, small, solid lesions. B. Gross pathology of Mtb-AG infected rabbit lungs showing multiple, large, solid lesions. C. Representative H&E stained lung section of Mtb-SC infected rabbits showing a large granuloma with immune cell accumulation and inflammatory foci, (*) surrounded by macrophages (m) and lymphocyte (ly) cuff. D. Representative H&E stained lung section of Mtb-AG infected rabbits showing a granuloma with large necrotic foci surrounded by neutrophils (arrows) and foamy macrophages (fm). E-F. Immunofluorescence imaging of Mtb (arrows) in Mtb-SC (E) and Mtb-AG (F) infected rabbit lung sections. Original magnification: 400x (C, D), 630x (E, F). G. Number of subpleural lung lesions in rabbit lungs infected with Mtb-AG or Mtb-SC; n= 6 per group. p-value was calculated using Mann-Whitney U-Test. H. Morphometric measurement of granulomatous lesion size in rabbit lungs infected with Mtb-SC or Mtb-AG. Data compiled from 24-30 fields per animal; n=4 per group per time point. p-value was calculated using Mann-Whitney U-Test. I. Pathological disease scoring of rabbit lungs infected with Mtb-SC or Mtb-AG. Data obtained from n=6 per group. *p<0.05; Mann-Whitney U-test.

The differential immune cell distribution in Mtb-AG or Mtb-SC infected rabbit lungs were measured by flow cytometry of single-cell suspension (Figure 3A and C) and immunohistochemistry (Figure 3B and D; Supplementary figure 3). A similar number of viable CD11b+ innate immune cells, including monocytes, neutrophils, and natural killer (NK) cells, were observed in Mtb-AG and Mtb-SC infected lungs at two weeks post-infection. However, a significantly higher number of CD11b+ cells was noted in the Mtb-SC infected animals at four weeks. (Figure 3A). In contrast, a significantly higher number of viable activated macrophages (IBA1+) was present in the Mtb-AG infected lesions at two and four weeks post-infection (Figure 3B). No significant difference in the number of CD4+ T-cells was observed between the Mtb-AG and Mtb-SC infected animals (Figure 3C), while a significantly higher number of CD8+ T-cells were noted in the Mtb-SC infected rabbit lungs at two and four weeks post-infection (Figure 3D).

**Figure 3.**
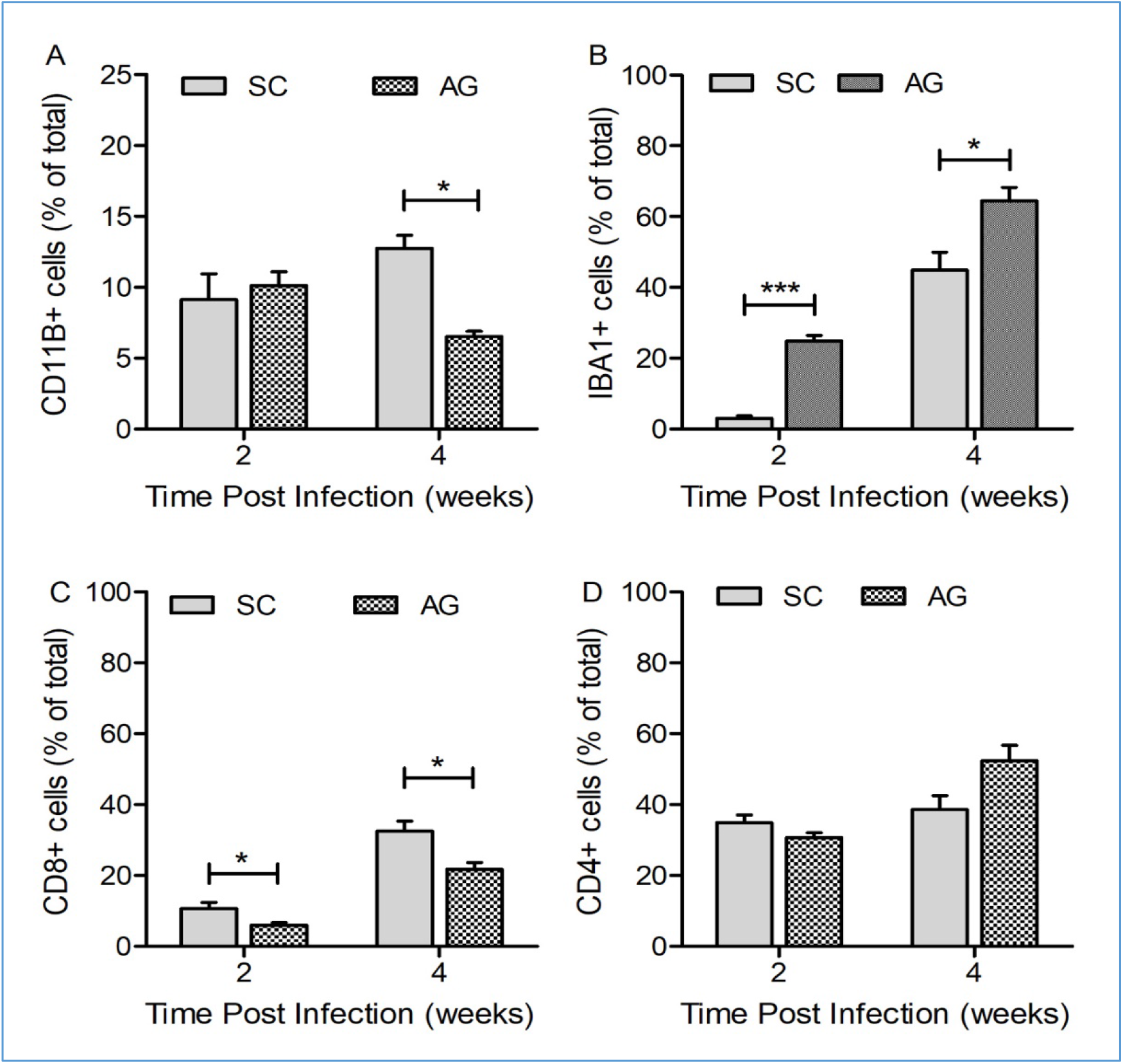
Immune cell distribution in the lungs after Mtb-AG or SC infection. Lung cells (A and C) or sections (B and D) from AG or SC infected rabbits at 2 and 4 weeks were stained specific antibodies against A. CD11b, B. IBA1 (macrophages), C. CD4 and D. CD8. Sections were analyzed microscopically, and cells positive for the markers were counted manually. Forty random fields from each animal were counted. n=3 animals Per group. Values plotted are mean +/− standard error. Data were analyzed by Mann-Whitney U-test. *p<0.05, **p<0.01, ***p<0.005.

### Mtb-AG exposure upregulates necrosis and cell death networks early during lung infection

To evaluate the early (i.e., 24 hours) differential host response in the lungs of rabbits exposed to Mtb-AG and Mtb-SC, RNAseq analysis was performed. The RNAseq data obtained were consistent within samples and reproducible across groups (Supplementary Figures 4–6). As shown in Figure 4, a higher number of genes were significantly perturbed in the Mtb-AG than Mtb-SC-infected rabbit lungs (1,552 versus 1,373 SDEG), with 822 genes commonly expressed between these two groups (Figure 4A and 4B). The pathway/network analysis of significantly differentially expressed genes (SDEG) revealed that inflammatory response (Figure 4C), host cell death and organismal morbidity (Figure 4D), and host cell necrosis (Figure 4F) were upregulated, while host cell survival (Figure 4F) was downregulated during Mtb-AG infection. Importantly, the gene expression pattern in Mtb-AG infected rabbit lungs revealed induction of apoptosis and necrosis and dampening of autophagy pathways (Supplementary Table 1). Other significantly differently regulated networks included host cell viability, cell survival, molecular export, autophagy, and cellular homeostasis (Table 1). Consistent with the perturbations in these biological functions, several canonical pathways associated with pro-inflammatory responses, including macropinocytosis signaling, GM-CSF signaling, dendritic cell maturation, and Th1 responses, were significantly downregulated during Mtb-AG infection. In contrast, genes involved in PTEN, PPAR, calcium, and CD27 signaling in immune cells were upregulated in Mtb-AG infected lungs (Table 2). Thus, the RNAseq analysis revealed early and significantly elevated host inflammatory responses and necrotic cell death pathways in Mtb-AG infected rabbit lungs.

**Figure 4.**
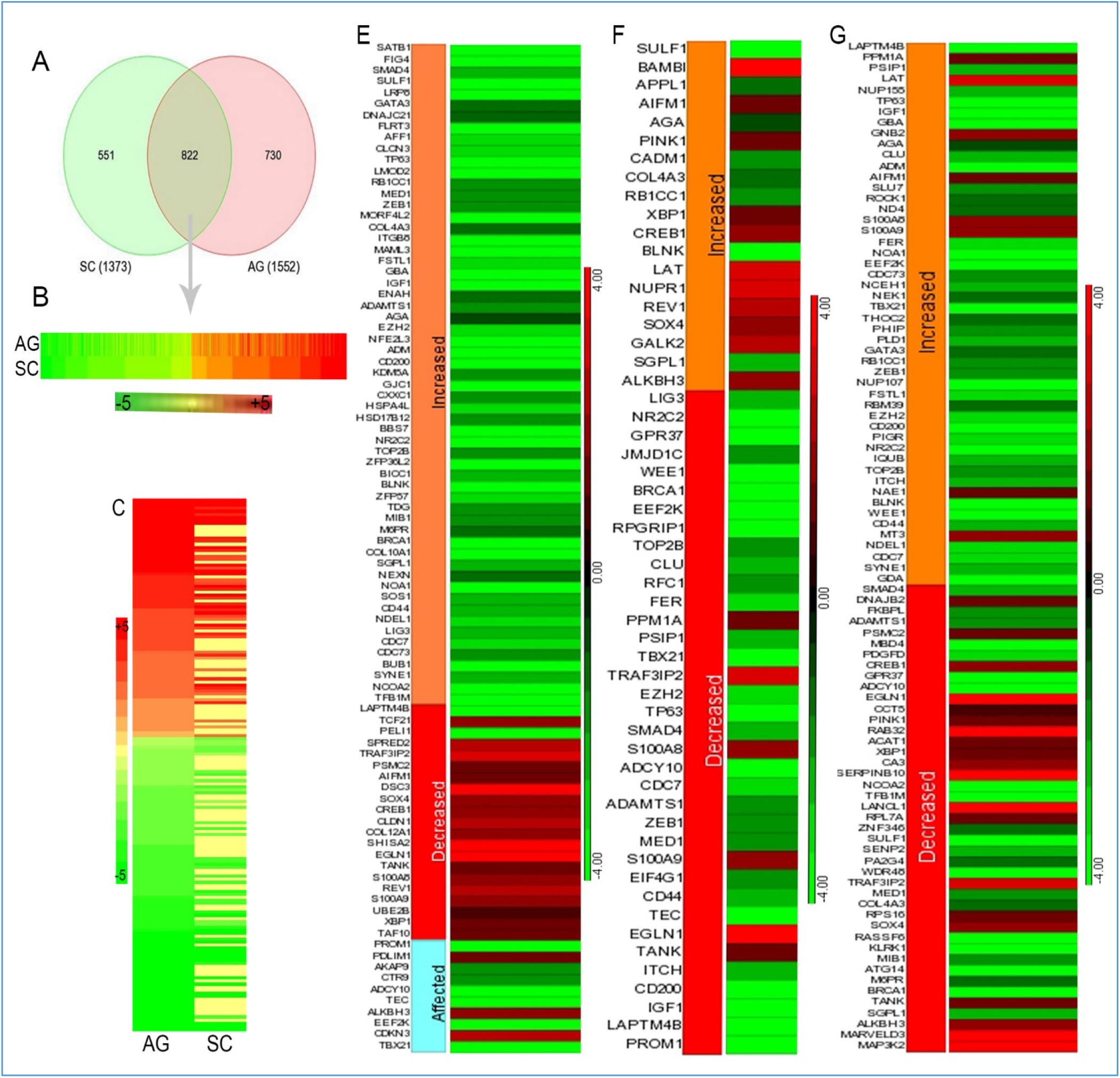
RNAseq analysis of host gene expression in rabbit lungs during Mtb-AG and Mtb-SC infection. A. Venn diagram showing significantly differentially expressed genes (SDEG) in Mtb-SC or –AG infected rabbit lungs at 24 hours (n=3 animals/group). Uninfected samples were used for normalization of gene expression from Mtb-infected samples. B. Heat map of SDEGs commonly perturbed during both Mtb-SC and -AG infection. Red color represents upregulation and green color represents downregulation of gene expression. The scale bar ranges from −5 (green) to +5 (red). C. Expression pattern of SDEGs showing upregulation of inflammatory response network in Mtb-AG. D. SDEGs expression showing upregulation of host cell death and organismal morbidity network in Mtb-AG, compared to Mtb-SC infection. E. SDEGs expression showing dampening of host cell survival network in Mtb-AG, compared to Mtb-SC infection. F. SDEGs expression showing increased host cell necrosis network in Mtb-AG, compared to Mtb-SC infection. The scale bars in D-F ranges from-4 (green) to +4 (red).

**Table-1.** List of significantly affected biological functions in Mtb-AG infected rabbit lungs

| **Biological Functions Annotation** | **Adjusted p-Value<sup>1</sup>** | **Activation z-score<sup>2</sup>** | **# genes** | **% consistency and direction<sup>3</sup>** |
| --- | --- | --- | --- | --- |
| Cell viability | 0.0114 | -2.40 | 53 | 66 (-ve) |
| Cell survival | 0.00453 | -2.31 | 58 | 66 (-ve) |
| Molecular Export | 0.0027 | -2.27 | 16 | 50 (-ve) |
| Autophagy | 0.0128 | -2.26 | 20 | 82 (-ve) |
| Cellular homeostasis | 0.0126 | -2.11 | 62 | 44 (-ve) |
| DNA repair | 0.000409 | -2.0 | 19 | 17 (-ve) |
| Necrosis of cells | 0.00443 | 2.40 | 171 | 67 (+ve) |
| Morbidity or mortality | 0.00465 | 4.36 | 92 | 64 (+ve) |
| Organismal death | 0.00651 | 4.36 | 90 | 64 (+ve) |
| Cell death and organismal injury | 0.00651 | 4.4 | 52 | 48(+ve) |

<sup>1</sup>compared to Mtb-SC infection data. <sup>2</sup> z-score is a measure of significance for the directionality of a biological function/pathway. It is derived from the gene expression pattern. A positive z-score indicates activation/upregulation, and a negative z-score indicates suppression/downregulation of a biological function. <sup>3</sup>represents the percentage of total genes with expression consistent with the activation/upregulation (+ve) or suppression/downregulation (-ve) of a biological network.

**Table 2.** List of up and downregulated canonical host signaling pathways in Mtb-AG infected rabbit lungs

| **Canonical pathways** | **z-score (AG-vs-SC)<sup>1</sup>** |
| --- | --- |
| PTEN signaling | 2.79 |
| RhoGDI signaling | 2.65 |
| PPAR signaling | 2.42 |
| Calcium signaling | 2.40 |
| CD27 signaling in lymphocytes | 2.36 |
| GAS signaling | 1.70 |
| p38MAPK signaling | 1.52 |
| Macropinocytosis signaling | -1.89 |
| ERK/MAPK signaling | -2.06 |
| Dendritic cell maturation | -3.01 |
| VEGF signaling | -3.05 |
| FcgRIIB signaling in B cells | -3.72 |
| GM-CSF Signaling | -4.22 |
| Th1 Pathway | -5.24 |

<sup>1</sup> z-score is a measure of significance for the directionality of a biological function/pathway derived from the gene expression pattern. A positive z-score indicates activation/upregulation, and a negative z-score indicates suppression/downregulation of a biological function.

### Mtb-AG induces early host inflammatory gene expression in the lungs

Next, we investigated the expression pattern of early inflammatory response markers in the lungs during Mtb-AG or Mtb-SC infection using qPCR. As shown in Figure 5, infection with Mtb-AG significantly upregulated the expression of the inflammatory molecules *S100A8*, *S100A9*, *S100A12*, *CRP*, *HIF1A*, *MMP1*, *MMP9*, *TBX21* and activated natural killer (NK) cell marker, *KLRG*, surface receptors, *PPARG*, *CD14*, and Th-2 type marker, *IL4* and *ARG1* (Figure 5). In contrast, expression of perforin (PRF1), cationic antimicrobial peptide (CAP18), and inflammasome (NLRP13) were upregulated in Mtb-SC infected lungs at 24hrs post-infection. These observations suggest that Mtb-AG induces a more robust host inflammatory response, while the antimicrobial response genes were preferentially upregulated in the Mtb-SC infected lungs.

**Figure-5.**
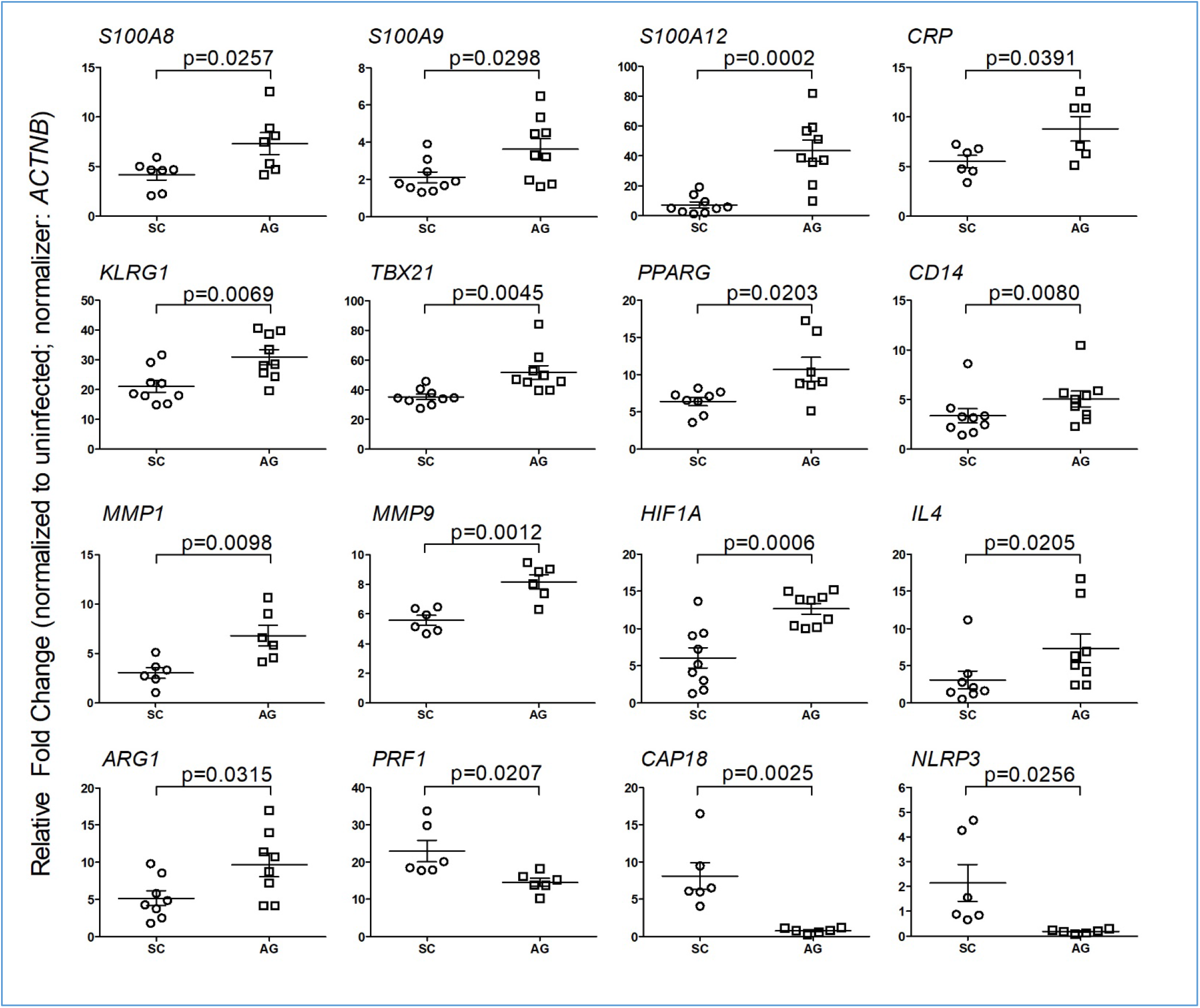
Modulation of host inflammatory response gene expression in Mtb-AG and Mtb-SC infected rabbit lungs. qPCR was used to measure the expression of selected host inflammatory response genes at 24 hours post-infection. The expression level of genes in uninfected lungs was used to normalize their corresponding levels in Mtb-AG and Mtb-SC infected samples. Beta-actin gene expression level was used as internal calibration control for each sample. The experiment was performed with n=3-4 per group per time point and repeated at least twice. p-value was calculated using Mann-Whitney U-Test.

### Mtb-AG induces cytotoxicity, cell death and dampens nitric oxide production in the infected lungs

Since Mtb-AG triggers a more robust inflammatory response and impacts activated immune cell distribution, we investigated the consequence of these processes on disease pathology by measuring the markers of host cell deaths in the rabbit lungs. Significantly increased LDH levels (a measure of cell death), the percentage of cytotoxicity and apoptotic cell death among total lung cells were noted in the Mtb-AG, compared to Mtb-SC infected animals as early as one week, and sustained until four weeks post-infection (Figure 6A-C). In contrast, nitric oxide (NO) levels were significantly higher in rabbit lungs during Mtb-SC-infection than Mtb-AG-infection at one week (Figure 6D). These data suggest that Mtb-AG infection is associated with elevated cytotoxicity and death of infected cells. Furthermore, the elevated and early production of antimicrobial NO2 during the early stages of infection might contribute to the control of Mtb-SC in the lungs. Consistent with this notion, Mtb-AG tolerated higher concentrations of agents that produce reactive oxygen and nitrogen species than Mtb-SC in broth cultures (Supplementary Figure 7).

**Figure-6.**
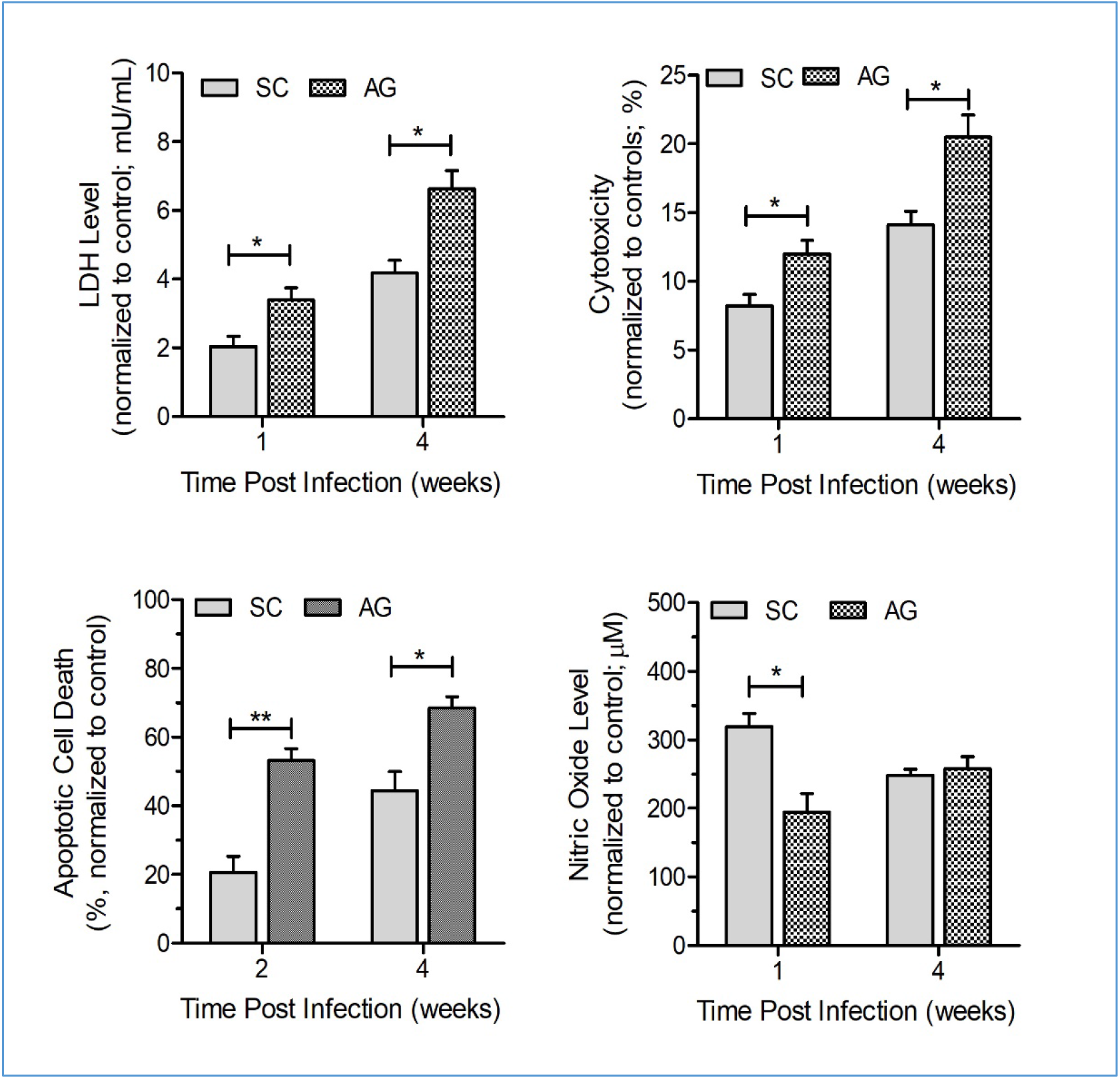
Mtb-AG exacerbates LDH levels, cytotoxicity, apoptosis, and dampens NO2 levels in the infected rabbit lungs. A. LDH levels were measured in lung homogenates and, B. cytotoxicity was calculated from the LDH levels of lung homogenates. C. Apoptotic cell death was calculated by TUNEL assay on rabbit lung sections at 2 and 4 weeks post-infection. D. Nitric oxide level was determined by Griess method using filtered cell lysate from infected rabbit lungs. All parameters were compared against un-infected control animals at 1 and 4 weeks post-infection. N=3 animals Per group. Values plotted are mean +/− standard error. Data were analyzed by Mann-Whitney U-test. *p<0.05, **p<0.01.

### Mtb-AG exacerbates host inflammatory responses and promotes tissue remodeling in the lungs

Since inflammation of the lung can impact its structure and function, we investigated the link between the inflammatory response and tissue remodeling that contribute to disease pathology by measuring the expression level of respective marker genes in the lungs infected with Mtb-AG or Mtb-SC. As shown in Table 3, infection with Mtb-AG significantly upregulated the expression of inflammatory molecules *S100A8*, *S100A9*, *CRP*, *IL23* activated natural killer cell marker, *KLRG*, and Th-2 type marker, *IL10* at one week post-infection (Table 3). A similar gene expression pattern was noted in Mtb-AG infected lungs by four weeks post-infection with a significantly higher level of *S100A8*, *S100A9*, *S100A12*, *IL17A*, *KLRG1*, *PPARG*, *ARG1* and *IL10*, and significantly dampened *TBX21* expression (Table 3). At the protein level, Mtb-AG induced more MMP9, IL-17 and IL-21, and lower TNF-α, IL-1α and NCAM levels than Mtb-SC at four weeks in the lungs, while IL-8, IL-1β, MIP-1β and leptin levels were similar between these two groups (Supplementary Figure 8). These observations suggest that induction of the host inflammatory response could contribute to the exacerbated cell death and disease pathology in Mtb-AG infected lungs.

**Table-3.**
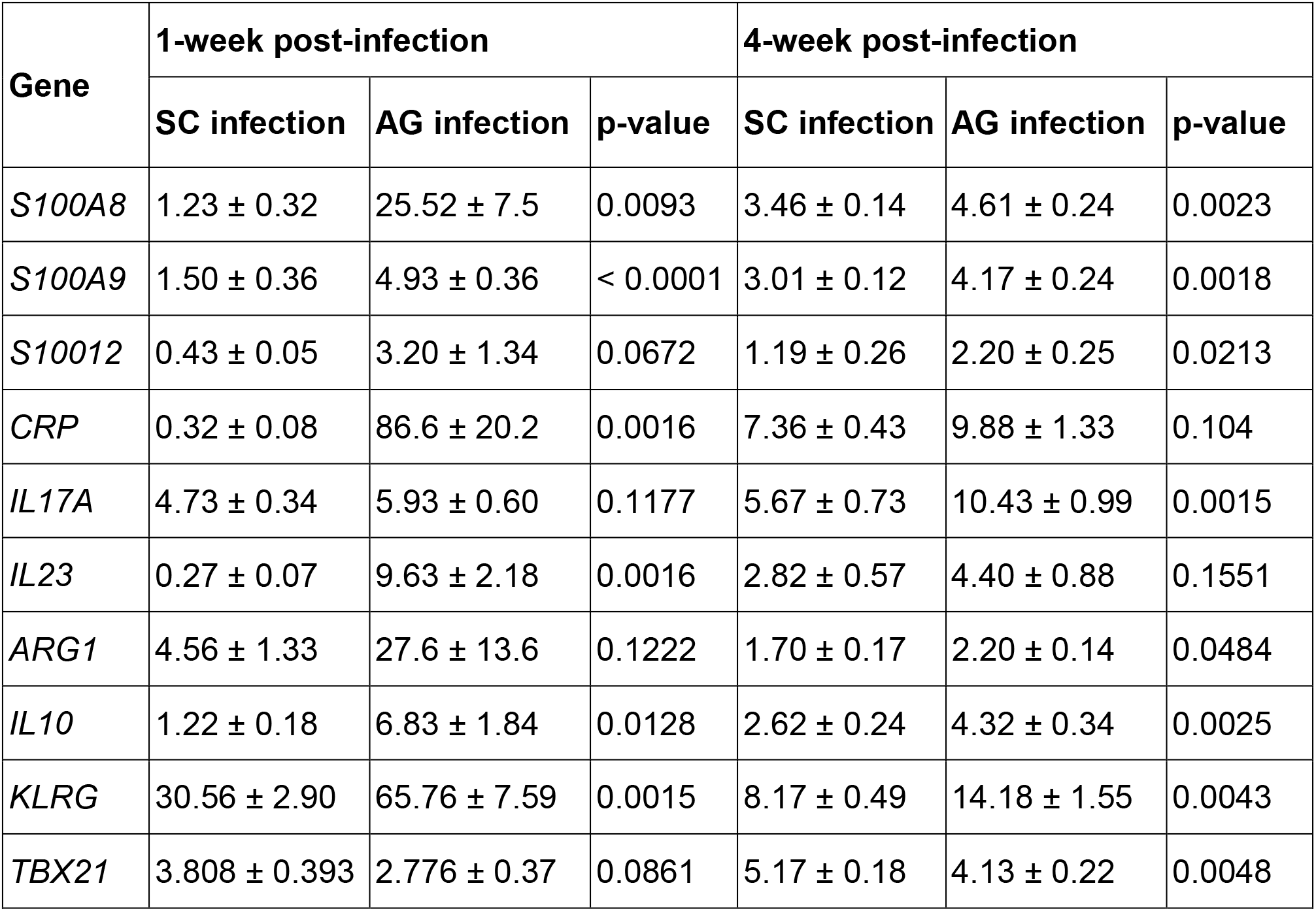
Expression level of host inflammatory response genes in rabbit lungs infected with Mtb-AG or Mtb–SC

| Gene | 1-week post-infection |  |  | 4-week post-infection |  |  |
| --- | --- | --- | --- | --- | --- | --- |
|  | SC infection | AG infection | p-value | SC infection | AG infection | p-value |
| *S100A8* | 1.23 ± 0.32 | 25.52 ± 7.5 | 0.0093 | 3.46 ± 0.14 | 4.61 ± 0.24 | 0.0023 |
| *S100A9* | 1.50 ± 0.36 | 4.93 ± 0.36 | < 0.0001 | 3.01 ± 0.12 | 4.17 ± 0.24 | 0.0018 |
| *S10012* | 0.43 ± 0.05 | 3.20 ± 1.34 | 0.0672 | 1.19 ± 0.26 | 2.20 ± 0.25 | 0.0213 |
| *CRP* | 0.32 ± 0.08 | 86.6 ± 20.2 | 0.0016 | 7.36 ± 0.43 | 9.88 ± 1.33 | 0.104 |
| *IL17A* | 4.73 ± 0.34 | 5.93 ± 0.60 | 0.1177 | 5.67 ± 0.73 | 10.43 ± 0.99 | 0.0015 |
| *IL23* | 0.27 ± 0.07 | 9.63 ± 2.18 | 0.0016 | 2.82 ± 0.57 | 4.40 ± 0.88 | 0.1551 |
| *ARG1* | 4.56 ± 1.33 | 27.6 ± 13.6 | 0.1222 | 1.70 ± 0.17 | 2.20 ± 0.14 | 0.0484 |
| *IL10* | 1.22 ± 0.18 | 6.83 ± 1.84 | 0.0128 | 2.62 ± 0.24 | 4.32 ± 0.34 | 0.0025 |
| *KLRG* | 30.56 ± 2.90 | 65.76 ± 7.59 | 0.0015 | 8.17 ± 0.49 | 14.18 ± 1.55 | 0.0043 |
| *TBX21* | 3.808 ± 0.393 | 2.776 ± 0.37 | 0.0861 | 5.17 ± 0.18 | 4.13 ± 0.22 | 0.0048 |

To determine the effect of Mtb-AG-induced inflammatory response on lung tissue remodeling, we investigated the expression of collagenases (MMP1, MMP13), stromelysin (MMP3), gelatinase (MMP9), elastase (MMP12), and membrane-type matrix metalloproteases (MMP14) by qPCR (Figure 7A). A significant increase in the expression of *MMP1*, *MMP3*, *MMP9*, *MMP12*, *MMP13* and *MMP14*, was noted in Mtb-AG infected, compared to Mtb-SC infected rabbit lungs, at both one and four weeks post-infection. Consistent with the upregulation of these MMPs, the Mtb-AG infected rabbit lungs showed more fibrin and collagen than Mtb-SC infected animals at four weeks post-infection, as revealed by the Masson’s trichrome staining (Figure 7B). These results suggest that active tissue damage/remodeling is associated with elevated MMP gene expression and inflammation mediated by host cell death in Mtb-AG infected rabbit lungs.

**Figure 7.**
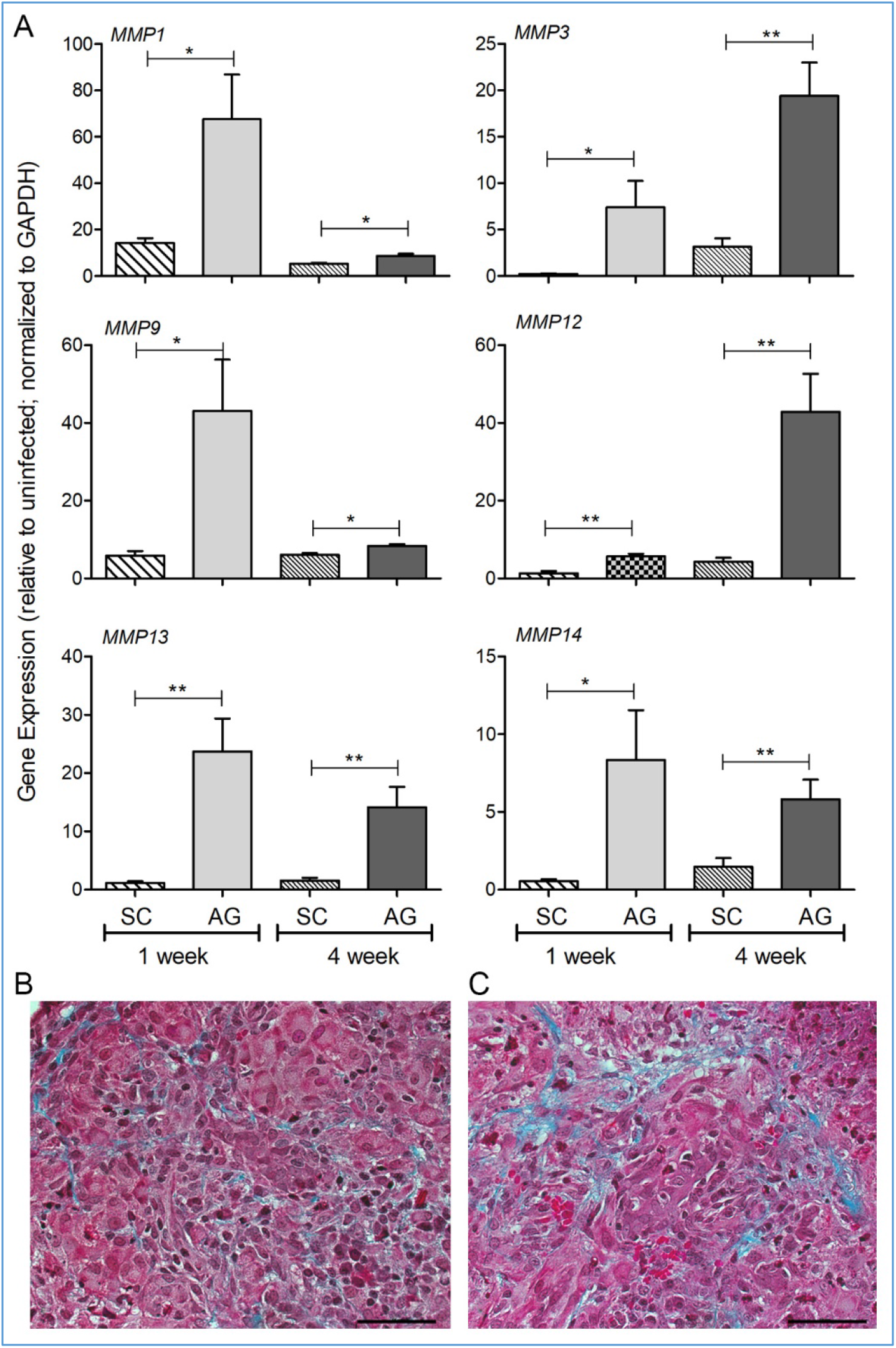
Mtb-AG induces more fibrosis and upregulates MMP gene expression in infected rabbit lungs. A. Expression level of MMP family genes was determined by qPCR using RNA from rabbit lungs infected with Mtb-AG or Mtb-SC at 1 and 4 weeks post-infection. Expression levels of each target gene were normalized to GAPDH levels. Levels in uninfected animals were used to calibrate the values in SC or AG-infected groups. The experiment was repeated with three biological replicates in duplicates. Values plotted are mean +/− standard error. Data were analyzed by Mann-Whitney U-test. *p<0.05, **p<0.01. B. Representative trichrome stained lung section of rabbits infected with Mtb-SC (left) or Mtb-AG (right), showing the extent of collagen deposition and fibrosis (blue color) at 4 weeks.

### Differential activation of T-cells in the spleen

Next, we evaluated the effect of Mtb-AG infection on the systemic immune response (i.e, spleen) by measuring the number of activated/proliferating CD4+ and CD8+ T-cells by flow cytometry. These T-cells from both Mtb-SC and Mtb-AG infected rabbits showed significantly higher proliferative response than uninfected controls at both two and four weeks post-infection (Figure 8A and B). However, the percent of activated/proliferating CD4+ and CD8+ T-cells in response to antigenic stimulation was significantly higher in Mtb-AG than Mtb-SC-infected animals at both two and four weeks post-infection. Together, these data suggest that compared to the lungs, a differential and robust immune activation occurs systemically in the spleen, during Mtb-AG, compared to Mtb-SC infection.

**Figure-8.**
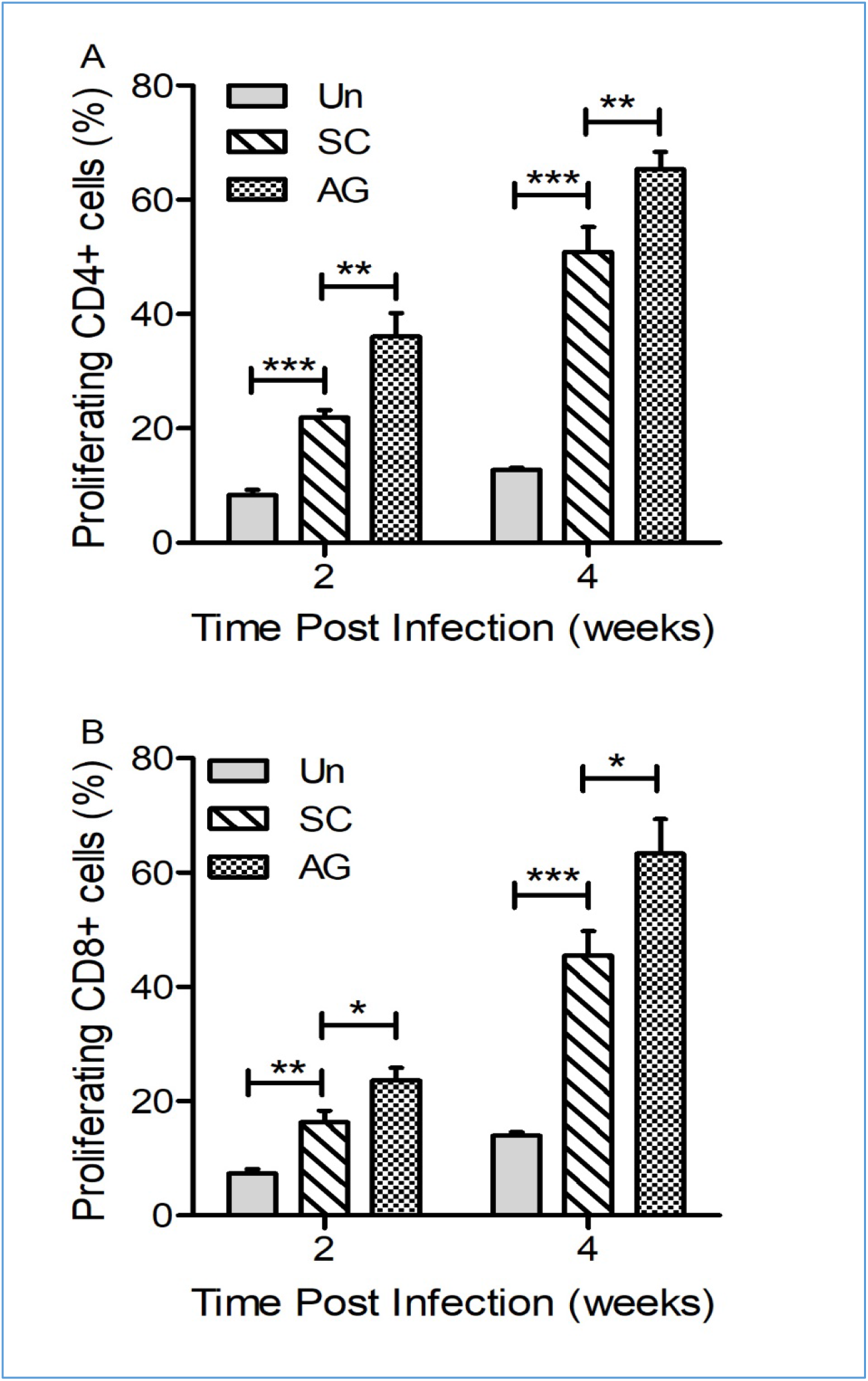
Activation of lung CD4+ and CD8+ T cells in Mtb-AG or SC infected rabbits. Percentage of proliferating CD4+ (A) and CD8+ (B) T-cells was determined at 2 and 4 weeks by flow cytometry analysis of single-cell suspensions using cell-specific antibodies after stimulation with heat-killed AG or SC or unstimulated. N=3 animals Per group. Values plotted are mean +/− standard error. Data were analyzed by one-way Anova with Tukey’s multi-group correction. *p<0.05, **p<0.01, ***p<0.005.

## DISCUSSION

We recently reported that in vitro infection of human monocyte-derived macrophages with Mtb-AG resulted in rapid phagocyte death, not seen with similar numbers of Mtb-SC. Additionally, efferocytosis of these cells by fresh macrophages caused rapid necrosis of the later cells, leading to a cascade of host cell destruction (10). Mtb-AG replicated efficiently in the infected/dead macrophages, while bacillary growth was limited in the Mtb-SC infected macrophages. In the present study, we tested the hypothesis that a similar differential response would be seen in vivo. That is, exposure of the rabbit lung to Mtb-AG would confer superior survival fitness on the bacilli in association with more extensive cell necrosis. We observed that Mtb-AG infection resulted in a sustained higher lung bacillary load, larger necrotic lesions, more severe inflammation, and extensive tissue remodeling in the lesions, compared to Mtb-SC infected animals. Consistently, several gene networks and pathways associated with necrotic cell death, cytotoxicity, and increased bacillary survival were upregulated early in Mtb-AG infected rabbit lungs.

Pathogen-mediated host cell death plays a crucial role in shaping the outcome of infection. Apoptosis, a programmed cell death process, and autophagy, a host-cell homeostasis mechanism, are implicated in controlling intracellular bacteria growth (13). Virulent Mtb has been shown to downregulate apoptosis and autophagy and promote host cell lysis by pyroptosis, necroptosis, and necrosis, thereby evading the antimicrobial responses of the host phagocytes (14, 15). Moreover, as we have shown in this study, dysregulation of apoptosis and autophagy can trigger inflammation and contribute to the progression of Mtb-induced pathology (16, 17)

The S100 family of proteins regulates the initiation and maintenance of inflammation in various disease conditions (18). S100A8, A9 and A12 are involved in neutrophil-mediated airway inflammation (19). These proteins were elevated during Mtb infection and shown to contribute to disease pathology in a murine model and in patients with active pulmonary tuberculosis (20,21,22). Consistent with these reports, we observed upregulation of S100A8, A9 and A12 expression in Mtb-AG infected rabbit lungs with high neutrophil infiltration.

The deficiency of KLRG1, a marker of NK cell activation, has been shown to enhance CD4+ T-cell responses, with increased production of IFN-γ and reduced bacterial burden in Mtb-infected mice (23). In addition, a reduction in Mtb load was associated with activated CD8+ T-cells secreting cytotoxic molecules, such as granulysin, perforin, and granzymes (24, 25). Moreover, limited activation of T-cells and suboptimal T-cell responses were reported in the granulomas of experimental animals and in human TB patients (26, 27, 28). Consistent with these reports, we observed a significant elevation in KLRG1 expression and dampened CD8+ T-cells activation in rabbit lungs. These results suggest that Mtb-AG can suppress the development of the anti-Mtb activity of NK cells and CD8+ T-cells. This may be due to the death of antigen-presenting phagocytes by Mtb-AG infection, leading to impaired adaptive immune responses. Additionally, the inflammation-driven cellular influx resulted in large, necrotic lesions in Mtb-AG infected rabbit lungs, in contrast to the primarily non-necrotic, smaller cellular granulomas seen in Mtb-SC infected lungs.

Infection with Mtb-AG leads to the upregulation of MMP genes and MMP9 protein, increasing fibrosis in the rabbit lungs. Several studies, including ours, have shown the association between elevated MMP gene expression and remodeling of the lung extracellular matrix (28–30). Moreover, MMP activity inhibition reduces morbidity and mortality of TB (31, 32) and inflammatory cytokines such as IL-17 and IL-23 regulate MMP expression and activity (33). In the present study, Mtb-AG infection was associated with increased ARG-1, IL-10, IL-17 and IL-23, potentially contributing to the induction of MMPs and fibrosis in the rabbit lungs.

Traditionally, both in vitro and in vivo experimental studies with Mtb have used deliberately dissociated aggregates/cording of the bacilli. Using tween in the growth medium and mechanical disruption, single-cell suspensions were generated, thereby facilitating an easier quantification of the bacilli used for infection and microbial characterization. Consequently, the potential role of Mtb-AG in driving pathogenesis has essentially been overlooked. Thus, the number of bacteria in the infectious inoculum also affects the outcome of the host response to Mtb. Thus, a higher initial bacterial load leads to more severe disease (6). In the current study, we show that the nature of Mtb implanted in the lungs (i.e., Mtb-AG versus Mtb-SC) can also impact the infection outcome. Thus, the various clinical outcomes of Mtb infection in humans, i.e., latent infection, active disease, and Mtb transmission, may be determined by the host response to the mechanical properties of Mtb initially implanted in the lungs. Future studies are warranted to characterize the nature of Mtb in infectious air droplets and understand their physiology in the context of TB epidemiology.

## METHODS

### Bacteria and chemicals

Pathogenic *Mycobacterium tuberculosis* H37Rv (Mtb) strain was obtained from Dr. Shinnick (US-CDC, Atlanta, GA, USA). Mtb-mCherry strain was obtained from Dr. Sigal. Cultures of Mtb-AG and Mtb-SC were prepared by growing bacteria with or without Tween-80, as described (10). De-clumping of AG was achieved by sonication (34). All chemicals were purchased from Sigma (Sigma-Aldrich, St. Louis, MO, USA) unless mentioned otherwise.

### Rabbit aerosol infection

Seventy-eight (n=78) female New Zealand white rabbits (*Oryctolagus cuniculus*) of about 2.5 kg body weight were purchased from Covance Inc (Covance Research Products, Denver, PA). Animals were randomly assigned into two groups and exposed to aerosols of Mtb H37Rv containing mostly Mtb-AG (n=39) or Mtb-SC (n=39) as described previously (34, 35). The rabbit aerosol infection chamber delivered bacterial inocula of mostly Mtb-AG and Mtb-SC as shown by staining of infectious aerosol particles collected at the delivery port directly on glass slides (Supplementary Figure 1). The infectious inoculum of Mtb-AG and Mtb-SC was adjusted to a similar number of total live bacteria as SC, based on the CFU from de-aggregated Mtb-AG. A positive correlation was observed between the lung CFU and Mtb fluorescence intensity (r^2^=0.7837) (Supplementary Figure 2A).

At 3hrs, 1-week, 2-weeks and 4-weeks post-infection, rabbits from each group (n=6-11 per group per time point) were euthanized, and lungs and spleen collected to evaluate bacterial CFU, histology, immune cell composition, enzyme levels, cellular function, and RNA isolation.

### Histology

Paraffin-embedded lung tissue was sectioned and stained with hematoxylin-eosin (H&E), acid-fast bacilli (AFB) or Masson’s trichrome staining as reported (35). Immunofluorescence with anti-Mtb primary antibody and Texas Red conjugated secondary antibody was used to visualize Mtb in lung sections as previously reported (36). The stained sections were analyzed in a Nikon Microphot DXM 1200C microscope and photographed using NIS-Elements software (Nikon Instruments Inc., Melville, NY, USA). Mtb fluorescent intensity was quantified using Image J (NIH, USA). Pathologoic analysis and scoring of lung sections were performed in a single-blinded way.

### Immunophenotyping of lung cells

Single-cell suspensions of rabbit lung cells were labeled with anti-CD4-Alexa647, anti-CD11b-PE (Biorad, Hercules, CA, USA) to enumerate CD4+ T-cells and innate immune cells, respectively, by using flow cytometry as described (34). Immuno-fluorescent staining with Anti-CD8-Alexa Flour 488 (Novus Biologicals, Centennial, CO, USA) and anti-IBA-1 (Abcam, MA, USA) was performed to determine the number of CD8+ T-cells and macrophages, respectively. Total and differential immune cells were counted in 20 fields per lung section per animal (n=3 per group per time point) using a Zeiss 200M fluorescent microscope (Carl Zeiss Microscopy LLC, White Plains, NY, USA).

### T-cell proliferation assay

Splenocytes were isolated from Mtb-AG and Mtb-SC infected rabbits (n=3 per group per time point) and labeled with carboxyfluorescein succinimidyl ester (CSFE) as described previously (37). Labeled cells were stimulated with heat-killed Mtb H37Rv or left unstimulated for 96 hrs, followed by staining with primary mouse anti-rabbit CD4 or anti-rabbit CD8 antibodies and secondary goat anti-mouse IgG-APC (ThermoFisher Scientific, CA, USA). Cell counts were measured and analyzed using BD Accurri C6 flow cytometer following the manufacturer’s instructions (BD Biosciences, CA, USA).

### Cytokine measurement

The levels of CCL4, CXCL8, MIP1B, IL1A, IL17A, IL1B, IL21, TNFA, MMP9, and NCAM-1 were measured in the rabbit lung homogenates (n=3 per group per time point) using Quantibody Rabbit Cytokine Array following manufacturer’s instructions (RayBiotech, Norcross, GA, USA).

### Quantification of Nitric Oxide

Nitrite concentration in the lung homogenate was measured (n=3 per group per time point) using Griess Reagent kit following manufacturer instructions (Sigma, St. Louis, MO, USA). Absorbance was recorded using a GloMax plate reader (Promega Corporation, Madison, WI, USA).

### Lactate dehydrogenase (LDH) activity, cytotoxicity, and apoptosis assays

LDH activity and percentage cell cytotoxicity in rabbit lungs (n=3 per group per time point) were determined using the LDH-Glow cytotoxicity kit as per the manufacturer’s instructions (Promega Corporation, Madison, WI, USA). Luminescence was recorded using a GloMax plate reader. Host cell apoptosis was measured in rabbit lung sections using the DeadEnd Colorimetric TUNEL System following the manufacturer’s instructions (Promega Corporation, Madison, WI, USA). Apoptotic and non-apoptotic cells were counted manually using the EVOS FL microscope (ThermoFisher Scientific, CA, USA).

### Lung RNA isolation

Total RNA from uninfected, Mtb-SC and Mtb-AG-infected rabbit lungs (n=3 per group per time point) was isolated using TRIzol reagent (Life Technologies, Grand Island, NY, USA) as described earlier (5, 38). The quality and quantity of the purified RNA were assessed by Bioanalyzer (Agilent Technologies Inc., Santa Clara, CA).

### RNAseq analysis

RNAseq experiment was performed using Lexogen QuantSeq 3′mRNA-Seq Library Prep Kit FWD for Illumina and NextSeq sequencing protocol by Lexogen (Lexogen, Inc, Greenland, NH, USA). Significantly differentially expressed genes (SDEG) were selected with a P<0.05 and 2-fold change in gene expression and used for network/pathway analysis using Ingenuity Pathway Analysis (IPA; Qiagen, Valencia, CA, USA) as described previously (5).

### Quantitative real-time PCR analysis (qPCR)

Total RNA isolated from Mtb-AG and Mtb-SC infected rabbit lungs (n=3 per group per time point) was used in qPCR experiments to measure the expression levels of *ARG1*, *TNFA*, *IL10*, *IL4*, *PPARG*, *S100A8*, *S100A9*, *S100A12*, *CRP*, *CD14*, *CD11B*, *NLRP3*, *CAP18*, *KLRG1*, *TBX21*, *HIF1A*, *ARG1*, *PRF*, *MMP1*, *MMP3*, *MMP9*, *MMP13*, *MMP14*, *IL10*, *IL17A* and *IL23* using Affinity QPCR cDNA Synthesis kit and Brilliant III Ultra-Fast SYBR® Green QPCR Master Mix as per kit instructions (Agilent Technologies Inc., Santa Clara, CA, USA).

### Mtb chromosome equivalent (CEQ) assay

Mtb genomic equivalent, a measure of total bacterial load in the rabbit lung (n=3 per group per time point), was determined by qPCR using Mtb16s rRNA gene and Brilliant III Ultra-Fast SYBR® Green QPCR Master Mix (Agilent Technologies, Inc. Santa Clara, CA, USA) as described earlier (39). A positive correlation was observed between the lung CFU and CEQ (r^2^=0.7881) (Supplementary Figure 2B).

### Mtb susceptibility test

Mtb-AG and Mtb-SC were grown in 7H9 media with different concentrations of hydrogen peroxide or sodium nitrite. Bacterial viability was determined by alamar blue assay as described previously (40).

### Statistical analysis

Mann-Whitney U-test and One-way Anova with Tukey’s correction were applied for pairwise and multiple group comparisons, respectively, using Prism-5 (GraphPad Software, La Jolla, CA, USA). A p< 0.05 was considered significant.

### Study approval

The Institutional Biosafety Committee and the Institutional Animal Care and Use Committee (IACUC) of Rutgers University approved all procedures involving rabbits. Animals were handled humanely according to the ethical guidelines of the federal Animal Welfare Act (AWA) and the Association for Assessment and Accreditation of Laboratory Animal Care International (AAALAC).

## AUTHOR CONTRIBUTIONS

SS and AS conceived the concept. SS, AS. GK designed the research studies, AK, RK, PS, AN and SS conducted experiments, acquired and analyzed data, AK, SS and GK wrote the manuscript. All authors have read, edited and agreed for submission.

## ACKNOWLEDGMENTS

SS and AS gratefully acknowledge the Bill and Melinda Gates Foundation for funding this study (# OPP1116944).

---

**Supplementary Figure-1.**
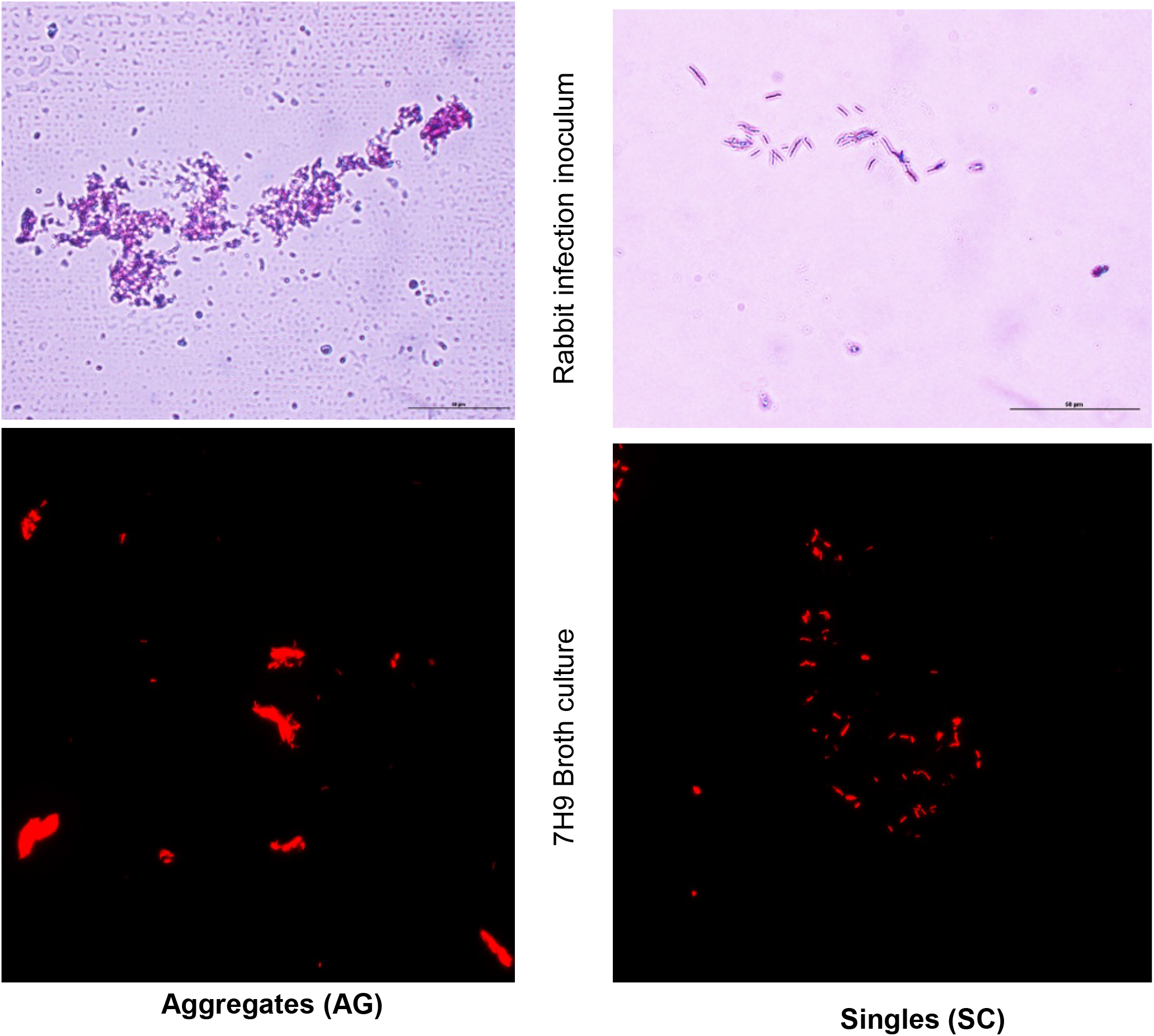
Morphology of Mtb in the Aerosols of Rabbit Infection Unit (C-H nose-only system) and in 7H9 broth culture. The top panel shows AFB-stained slides of infection inoculum captured at the delivery port of the rabbit infection chamber. Image captured at 630x magnification. Scale bar 10um. The bottom panel shows mCherry-expressing Mtb-H37Rv as aggregates (AG) or singles (SC) grown without or with tween-80, respectively. Image captured at 630x magnification.

**Supplementary Figure-2.**
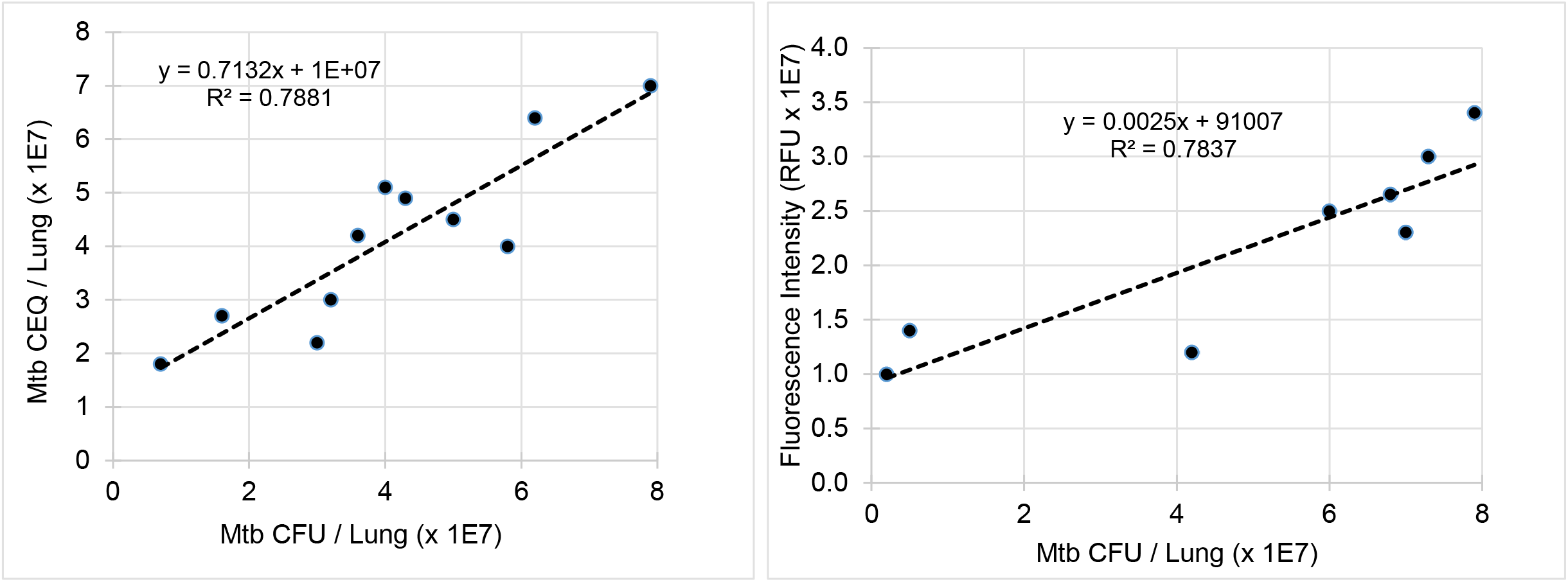
Correlation analysis between Mtb CFU, CEQ and bacterial fluorescent intensity. A. correlation between Mtb CFU and CEQ obtained from rabbit lungs at 4 weeks post-infection. B. correlation between Mtb CFU and bacterial fluorescence intensity measured in rabbit lungs at 4 weeks post-infection.

**Supplementary Figure-3.**
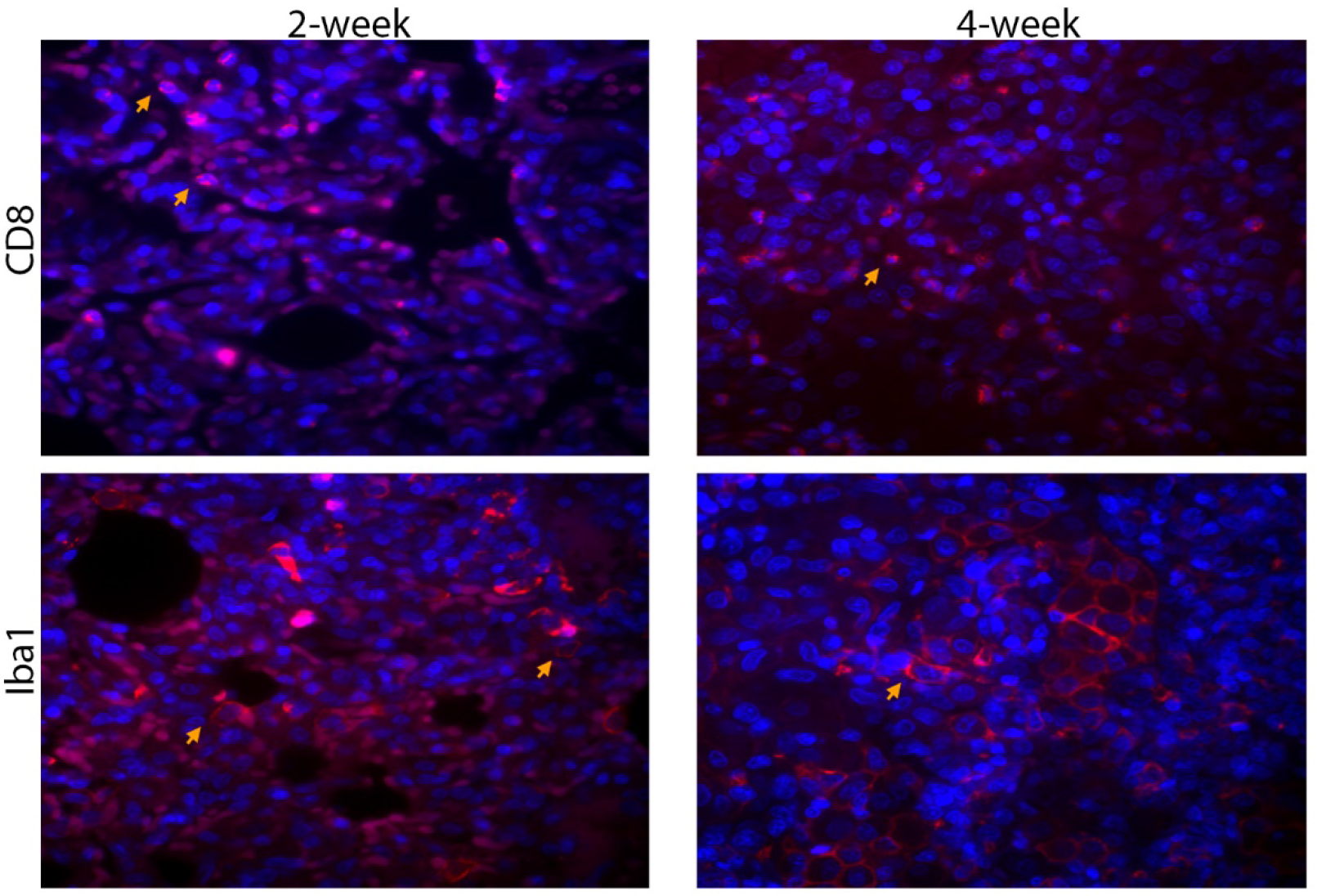
Representative immunofluorescent imaging showing CD8+ and Iba1+ (macrophages) in rabbit lung sections (red/pink spots; yellow arrows) infected with Mtb-AG or Mtb-SC at 4 weeks post infection. Nucleus is stained blue (DAPI).

**Supplementary Figure-4.**
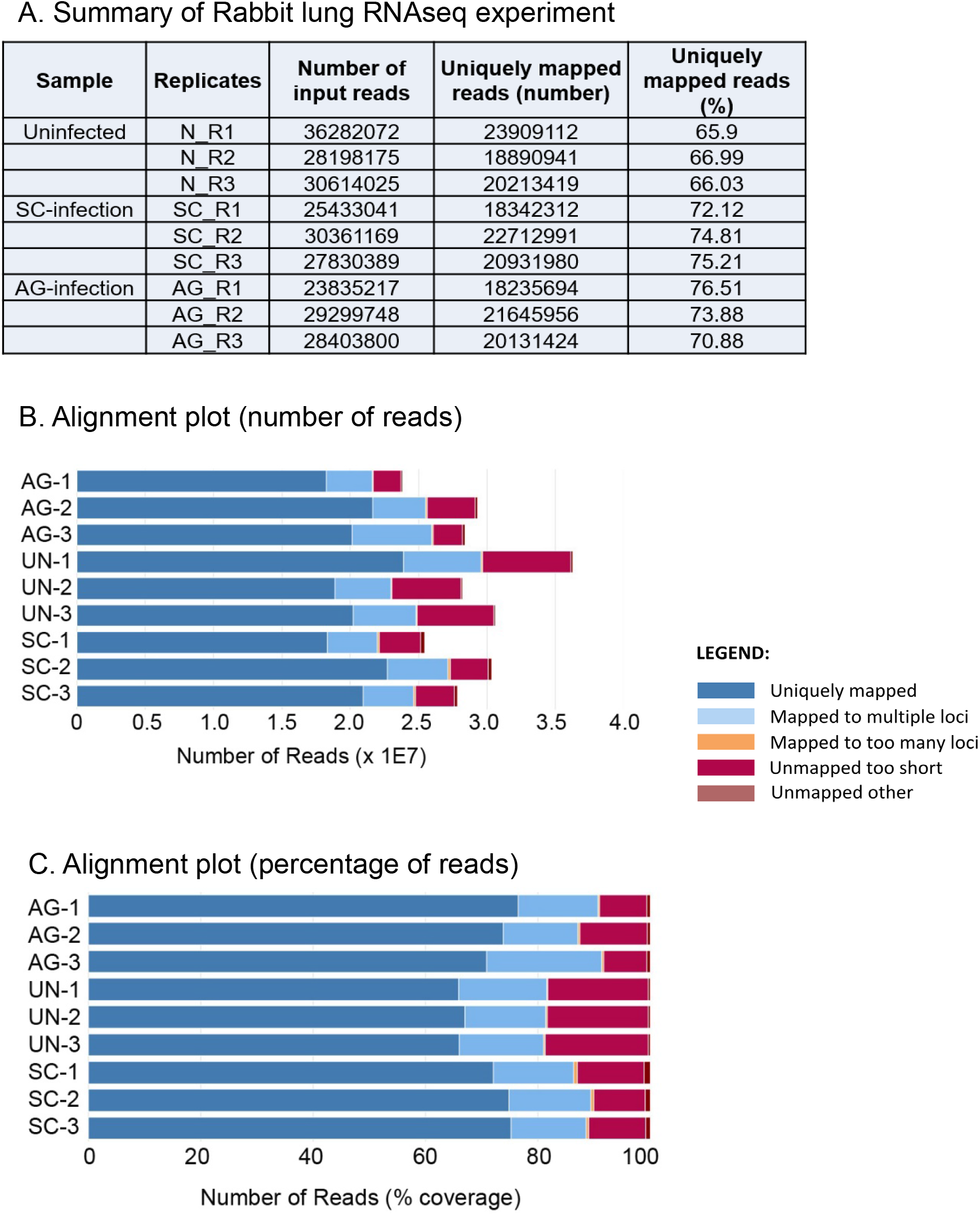
Summary of RNAseq data from uninfected or Mtb-AG or Mtb-SC infected rabbit lungs at 24 hours post-infection. A. Table showing the number of input reads and uniquely mapped reads. B. Alignment plot of RNAseq represented as the number of reads (x-axis) and sample type (y-axis). C. Alignment plot of RNAseq represented as % of reads (x-axis) and sample type (y-axis). The experiment was performed in triplicates for each sample type.

**Supplementary Figure 5.**
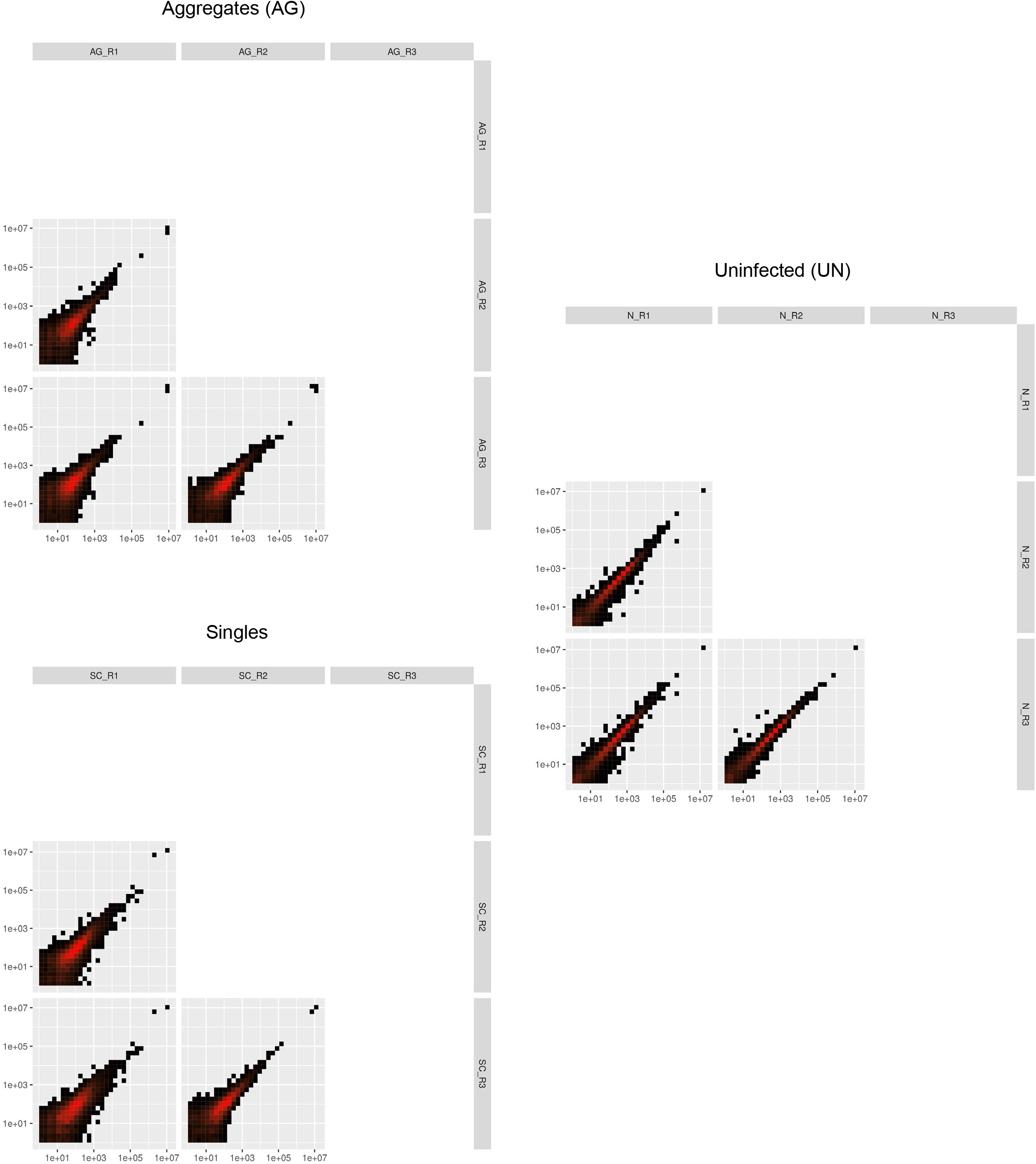
Reproducibility of biological replicates in RNAseq experiments. Plot shows significant consistency among the RNAseq reads obtained from triplicates of Mtb-AG or Mtb-SC infected or uninfected (UN) rabbit lung samples

**Supplementary Figure 6.**
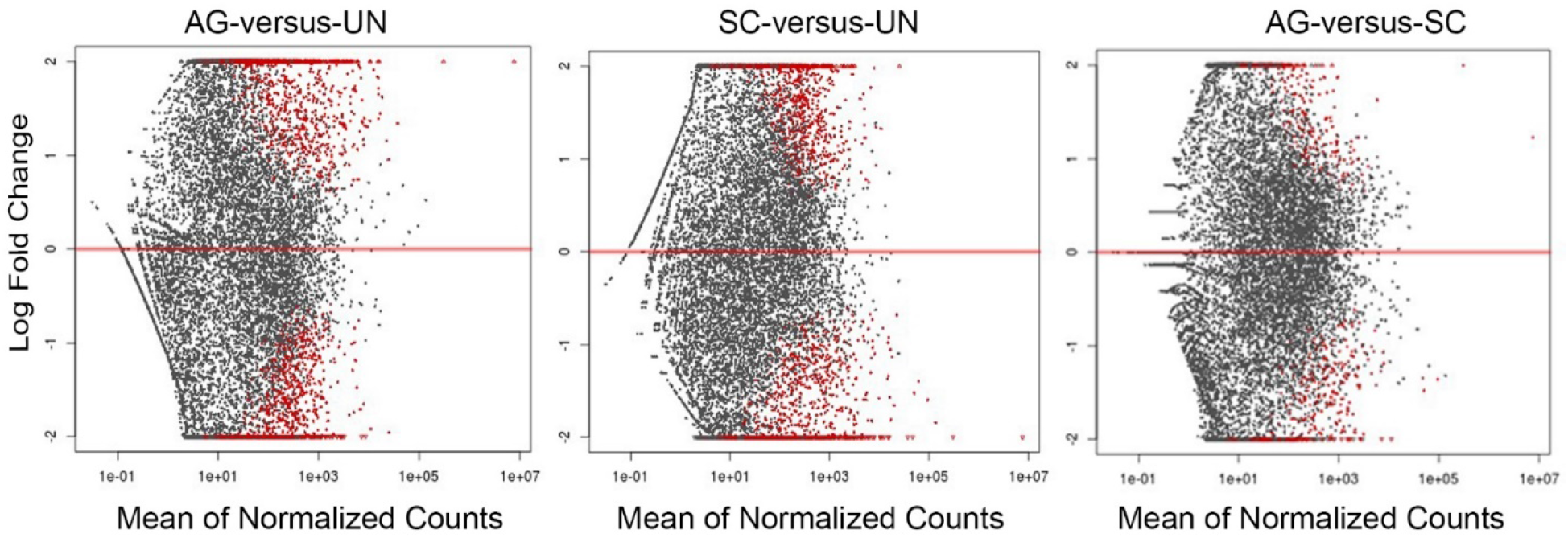
MA plot of normalized gene counts in Mtb-AG, Mtb-SC or uninfected (UN) rabbit lungs. Each dot is a gene count. Dots in red color are significantly differentially expressed in the comparator groups

**Supplementary Figure-7.**
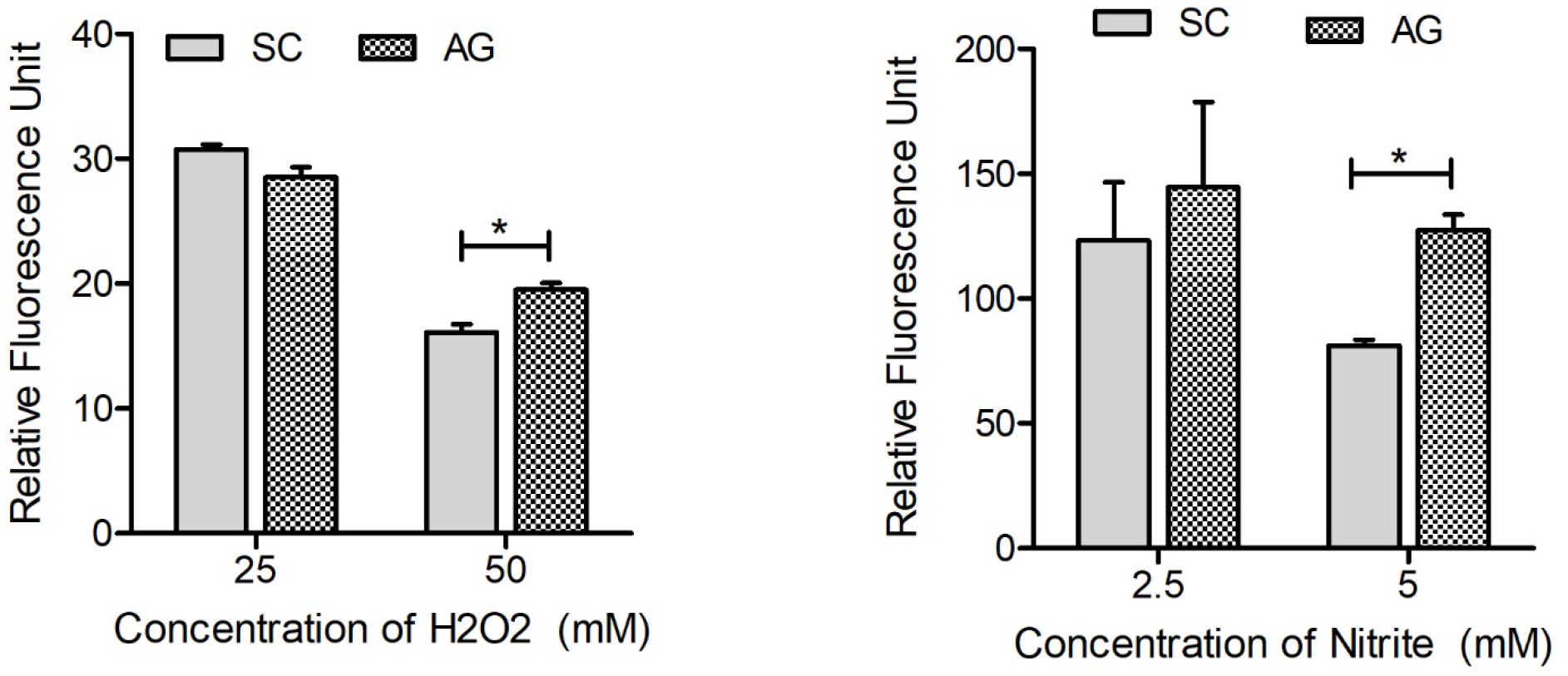
Mtb-AG is more resistant than Mtb-SC to exposure to ROS and RNS generating agents. Broth cultures of fluorescent Mtb AG and SC expressing mCherry were exposed to H2O2 or sodium nitrite, and bacterial viability was measured after 72 hours as fluorescent units. Untreated Mtb cultures were used to normalize the data from treated cultures. The experiment was repeated three times in duplicates. Values plotted are mean +/− standard error. Data were analyzed by Mann-Whitney U-test. *p<0.05.

**Supplementary Figure 8.**
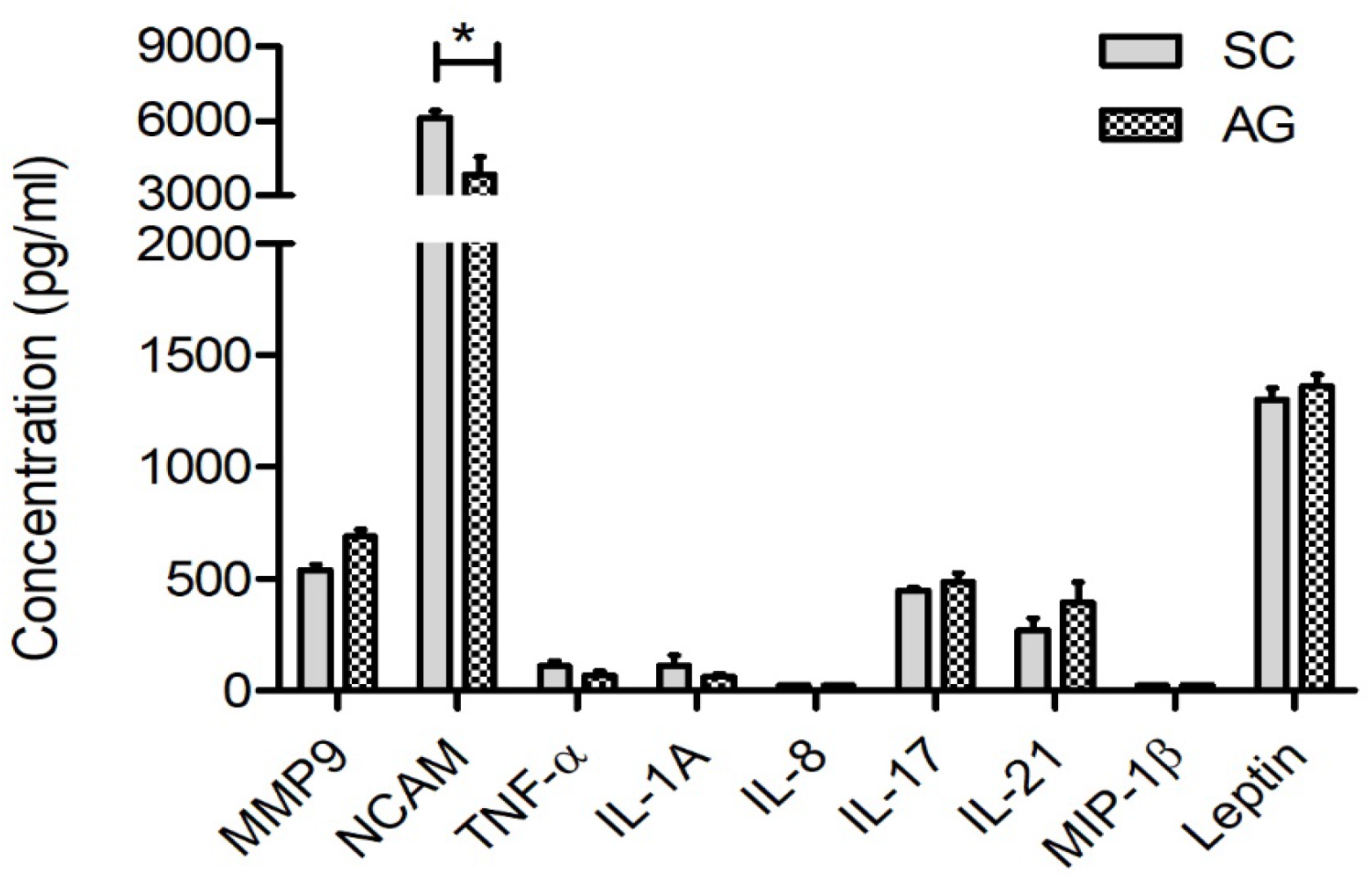
Levels of inflammatory cytokines/chemokines in rabbit lungs infected with Mtb-SC or AG at 4 weeks. Filtered lung homogenates were used to measure the markers by ELISA. n=3 animals Per group. Values plotted are mean +/− standard error. Data was analyzed by Mann-Whitney U-test. *p<0.05.

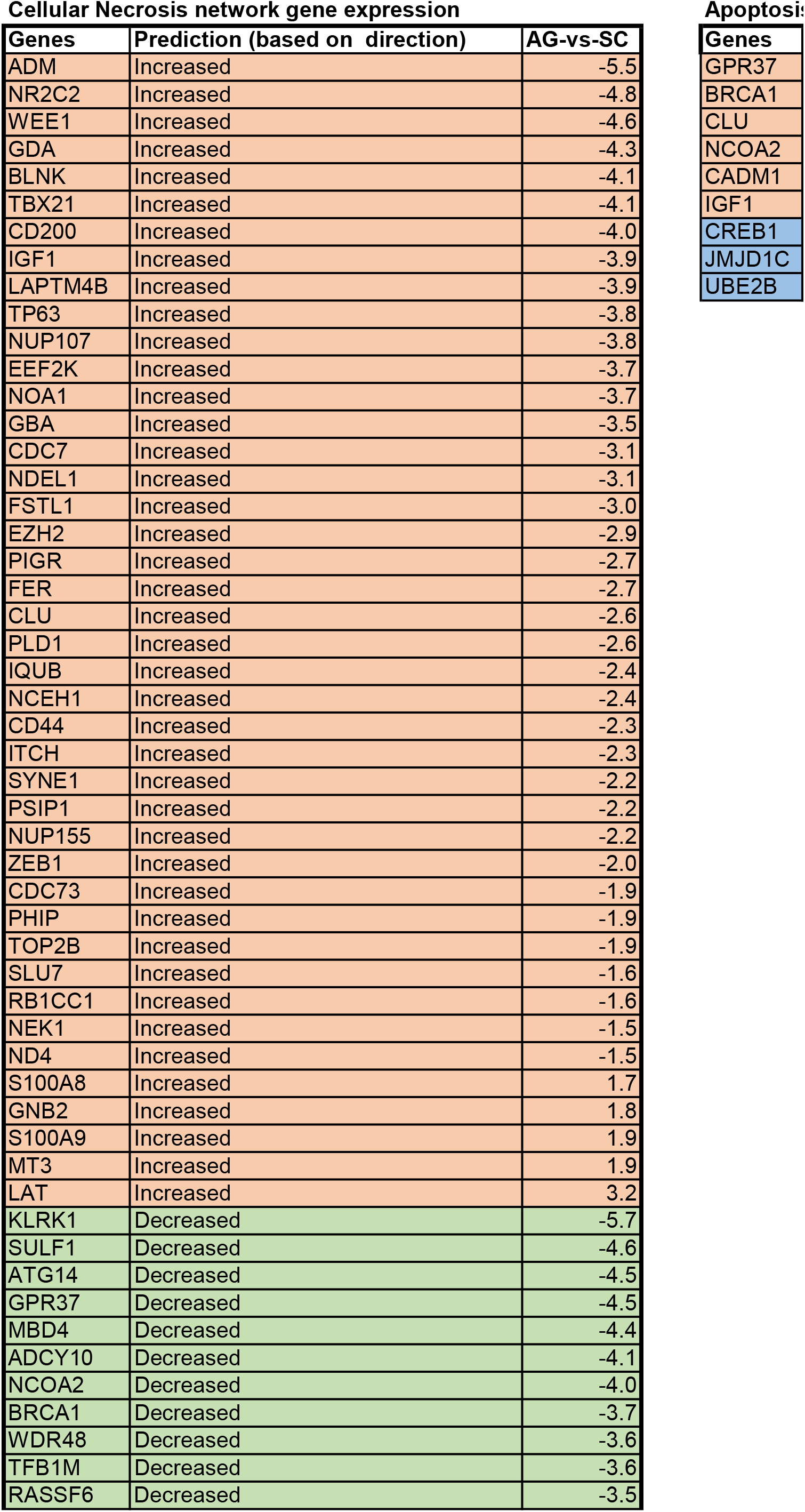
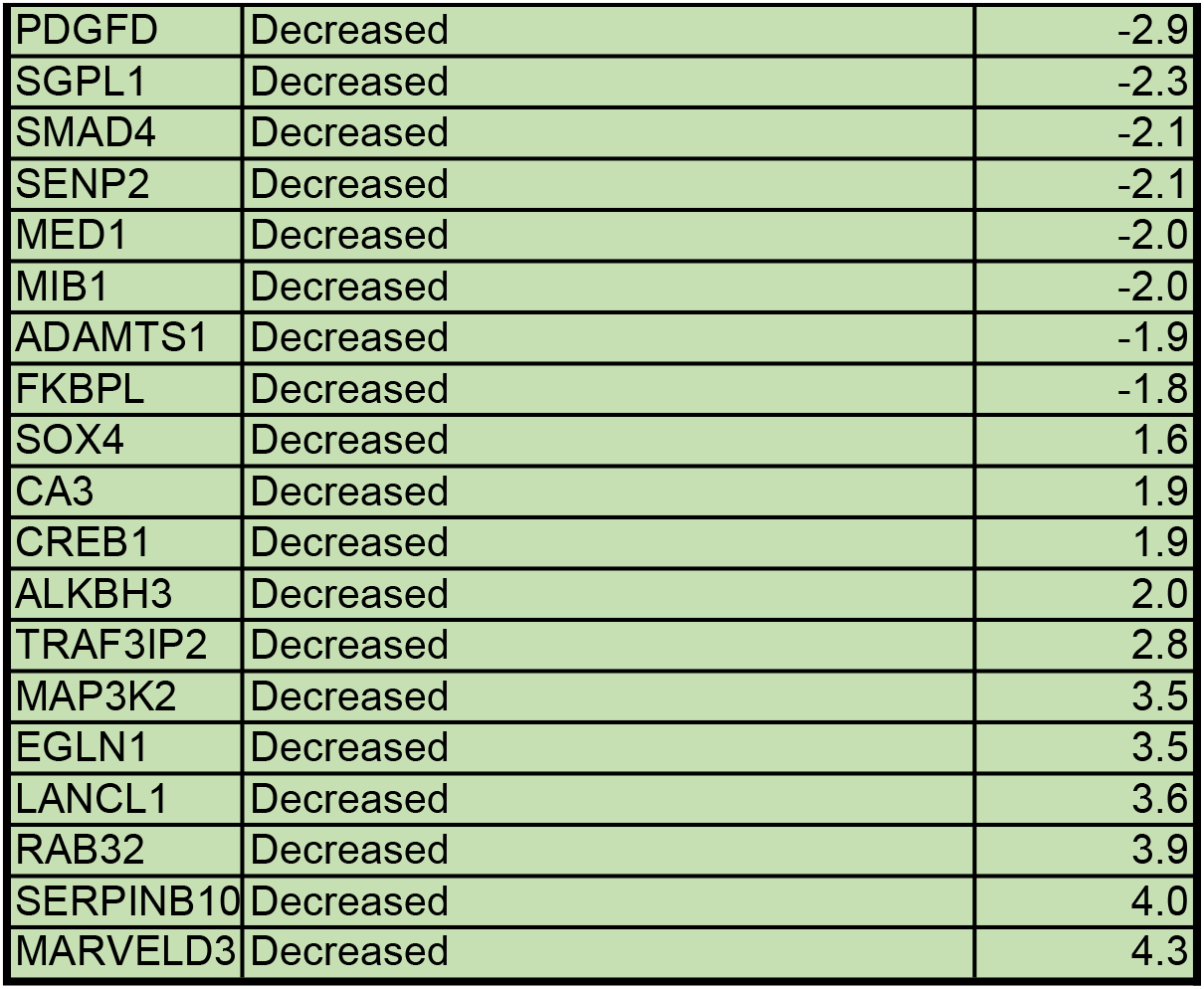
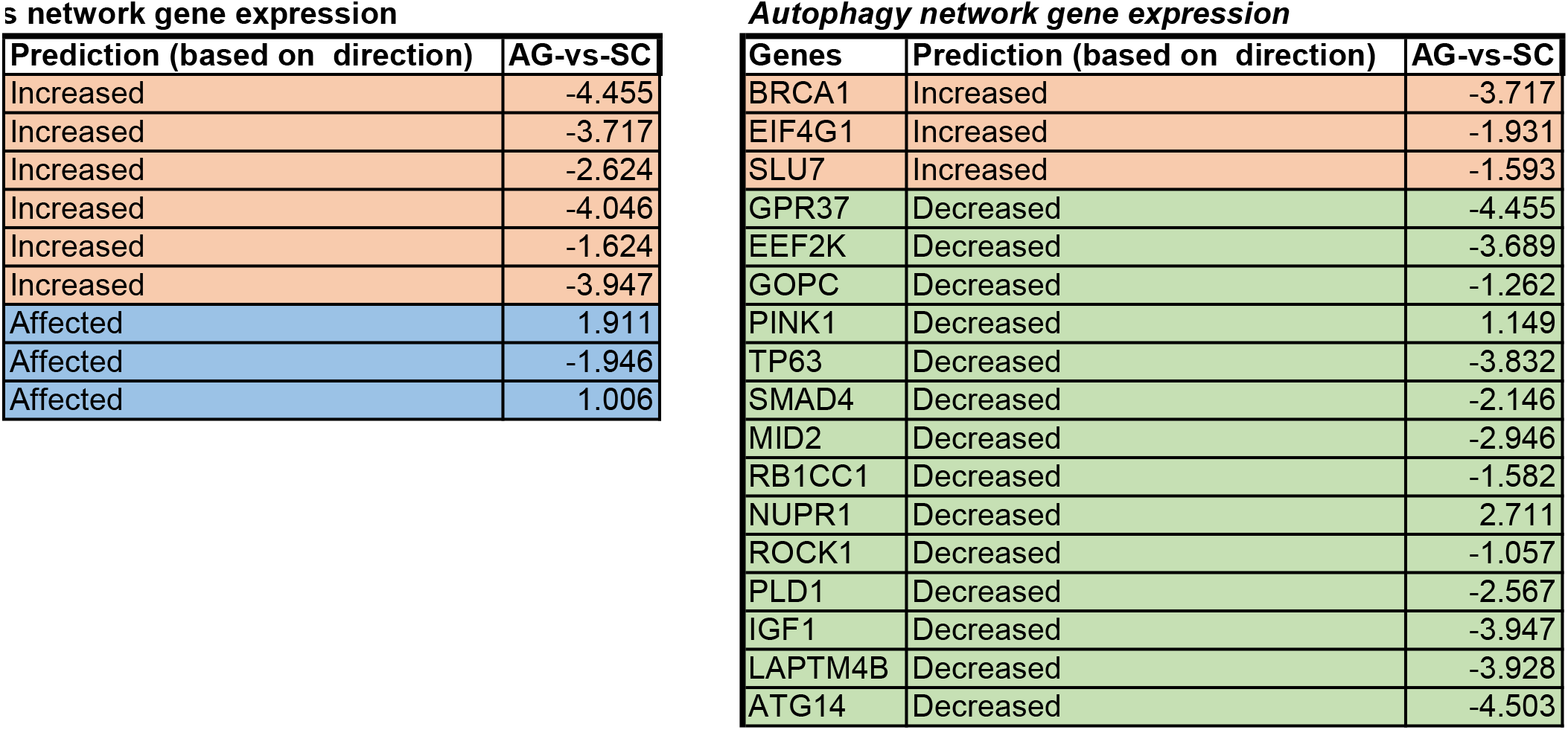

