## Supplemental files for "Aggregation state of *Mycobacterium tuberculosis* impacts host immunity and augments pulmonary disease pathology"

**SUPPLEMENTARY FIGURE-1**

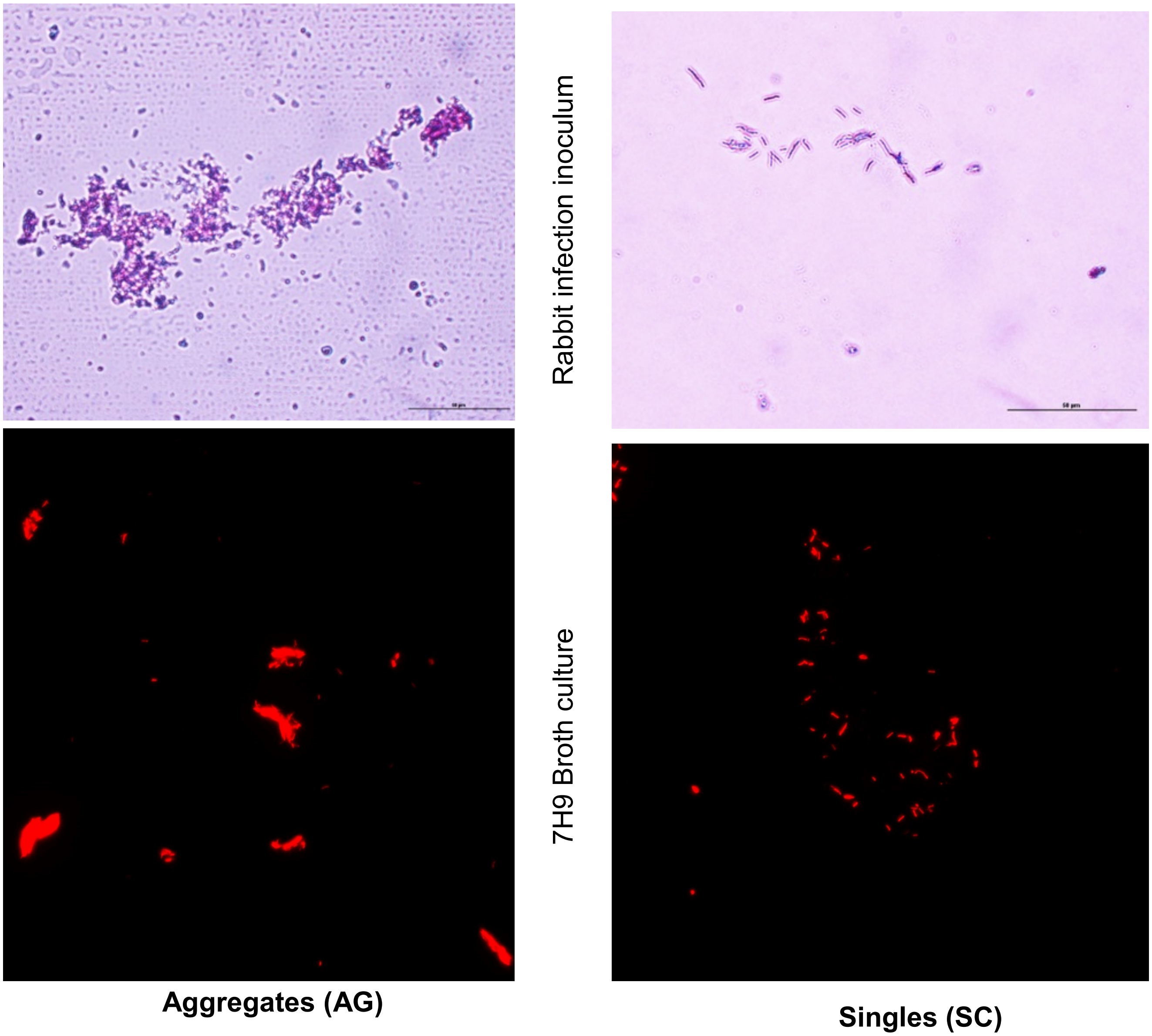


**Supplementary Figure-1.** Morphology of Mtb in the Aerosols of Rabbit Infection Unit (C-H nose-only system) and in 7H9 broth culture. The top panel shows AFB-stained slides of infection inoculum captured at the delivery port of the rabbit infection chamber. Image captured at 630x magnification. Scale bar 10um. The bottom panel shows mCherry-expressing Mtb-H37Rv as aggregates (AG) or singles (SC) grown without or with tween-80, respectively. Image captured at 630x magnification.

**SUPPLEMENTARY FIGURE-2**

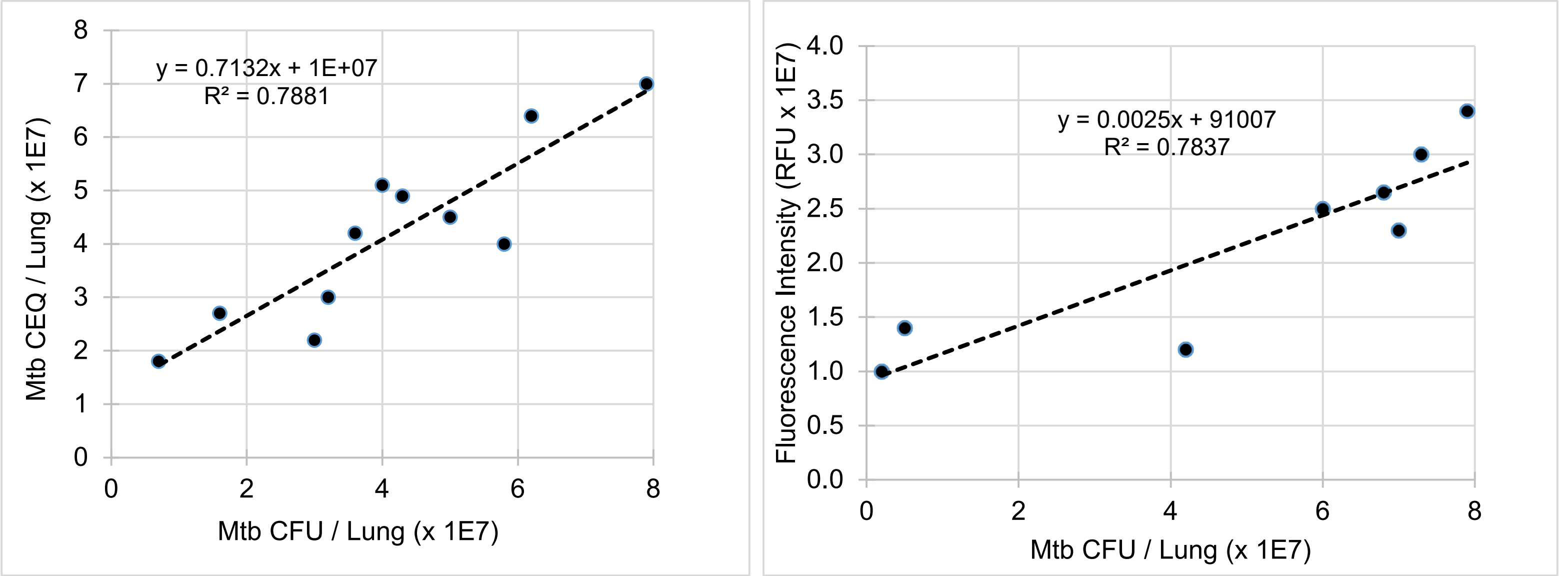


**Supplementary Figure-2.** Correlation analysis between Mtb CFU, CEQ and bacterial fluorescent intensity. A. correlation between Mtb CFU and CEQ obtained from rabbit lungs at 4 weeks post-infection. B. correlation between Mtb CFU and bacterial fluorescence intensity measured in rabbit lungs at 4 weeks post-infection.

**SUPPLEMENTARY FIGURE-3**

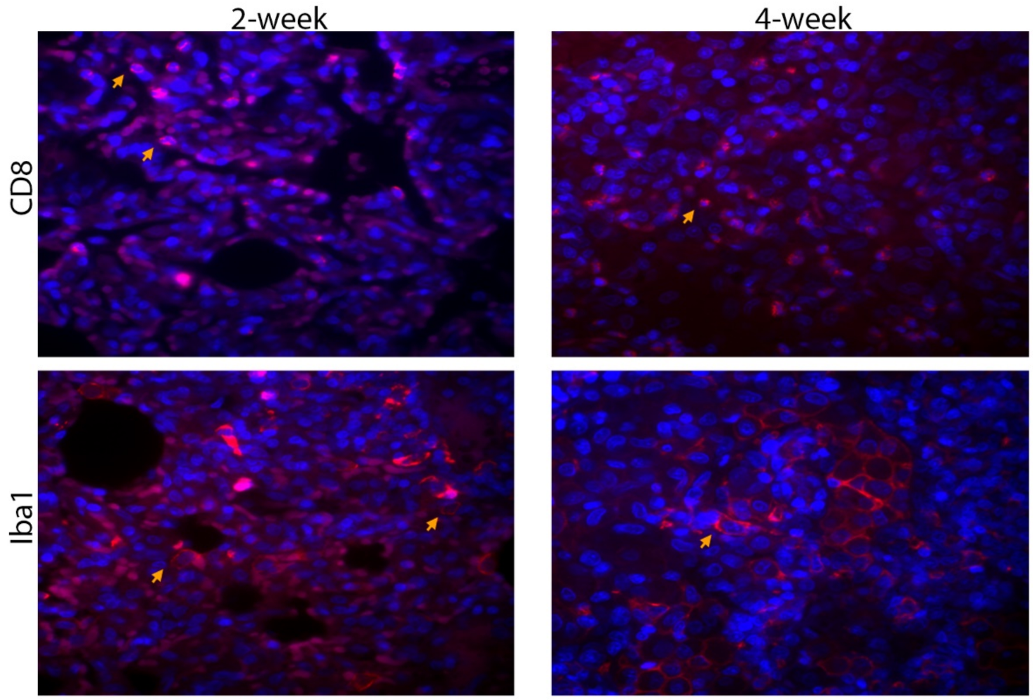


**Supplementary Figure-3**. Representative immunofluorescent imaging showing CD8+ and Iba1+ (macrophages) in rabbit lung sections (red/pink spots; yellow arrows) infected with Mtb-AG or Mtb-SC at 4 weeks post infection. Nucleus is stained blue (DAPI).

**SUPPLEMENTARY FIGURE-4**

A. Summary of Rabbit lung RNAseq experiment

| Sample | Replicates | Number of input reads | Uniquely mapped reads (number) | Uniquely mapped reads (%) |
|---|---|---|---|---|
| Uninfected | N_R1 | 36282072 | 23909112 | 65.9 |
| | N_R2 | 28198175 | 18890941 | 66.99 |
| | N_R3 | 30614025 | 20213419 | 66.03 |
| SC-infection | SC_R1 | 25433041 | 18342312 | 72.12 |
| | SC_R2 | 30361169 | 22712991 | 74.81 |
| | SC_R3 | 27830389 | 20931980 | 75.21 |
| AG-infection | AG_R1 | 23835217 | 18235694 | 76.51 |
| | AG_R2 | 29299748 | 21645956 | 73.88 |
| | AG_R3 | 28403800 | 20131424 | 70.88 |

B. Alignment plot (number of reads)

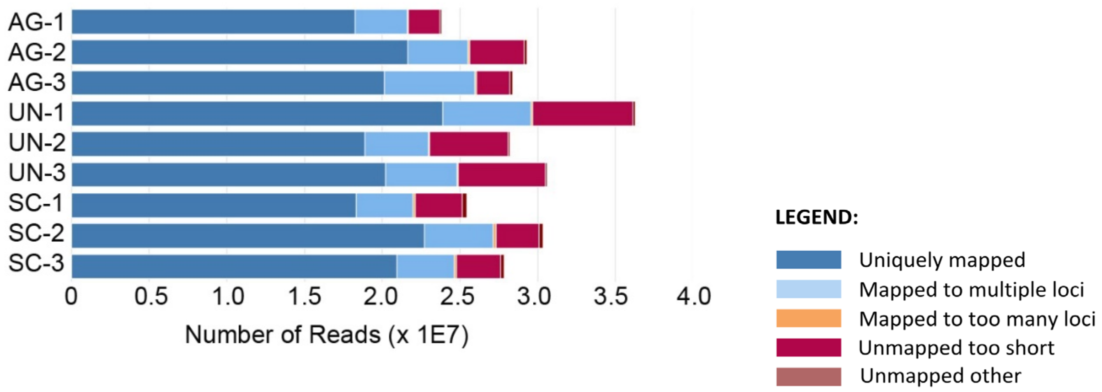


C. Alignment plot (percentage of reads)

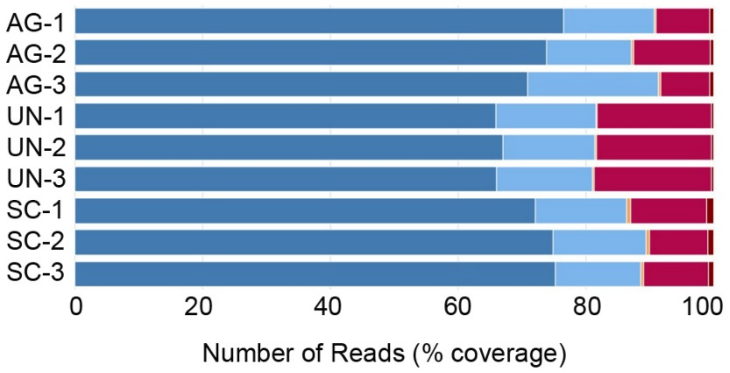


**Supplementary Figure-4**. Summary of RNAseq data from uninfected or Mtb-AG or Mtb-SC infected rabbit lungs at 24 hours post-infection. A. Table showing the number of input reads and uniquely mapped reads. B. Alignment plot of RNAseq represented as the number of reads (x-axis) and sample type (y-axis). C. Alignment plot of RNAseq represented as % of reads (x-axis) and sample type (y-axis). The experiment was performed in triplicates for each sample type.

**SUPPLEMENTARY FIGURE-5**

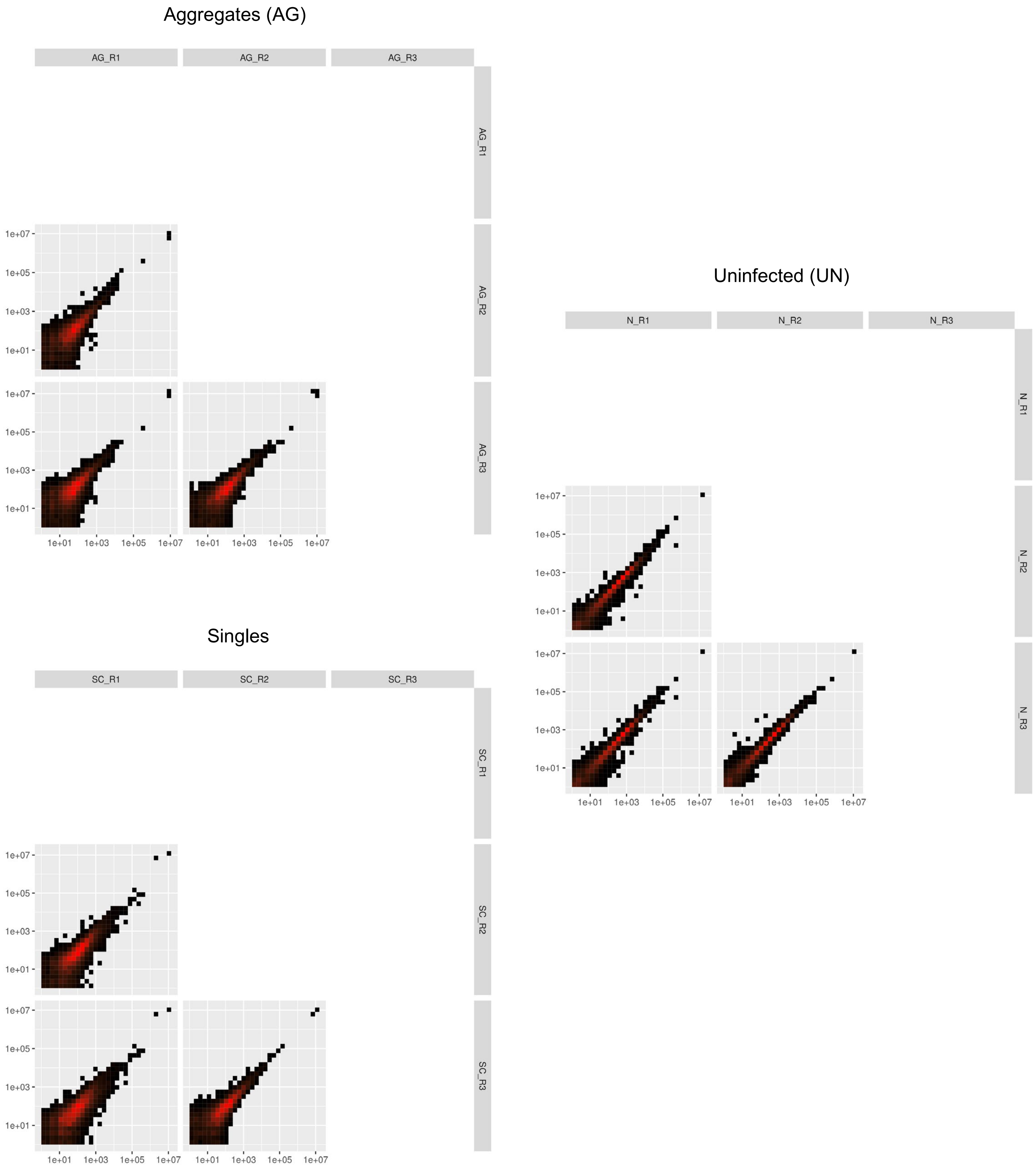


**Supplementary Figure 5.** Reproducibility of biological replicates in RNAseq experiments. Plot shows significant consistency among the RNAseq reads obtained from triplicates of Mtb-AG or Mtb-SC infected or uninfected (UN) rabbit lung samples

**SUPPLEMENTARY FIGURE-6**

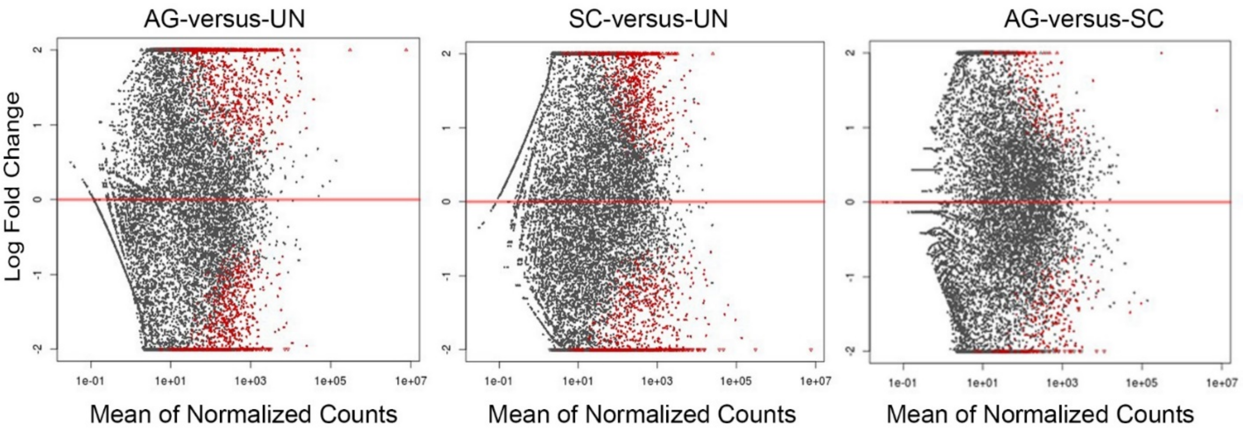


**Supplementary Figure 6.** MA plot of normalized gene counts in Mtb-AG, Mtb-SC or uninfected (UN) rabbit lungs. Each dot is a gene count. Dots in red color are significantly differentially expressed in the comparator groups

**SUPPLEMENTARY FIGURE-7**

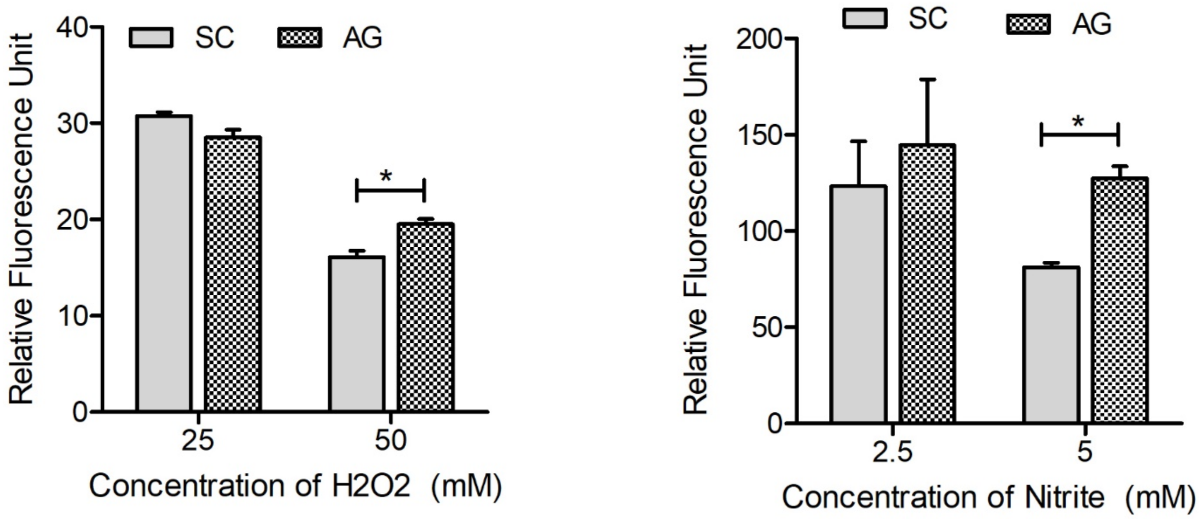


**Supplementary Figure-7.** Mtb-AG is more resistant than Mtb-SC to exposure to ROS and RNS generating agents. Broth cultures of fluorescent Mtb AG and SC expressing mCherry were exposed to $H_2O_2$ or sodium nitrite, and bacterial viability was measured after 72 hours as fluorescent units. Untreated Mtb cultures were used to normalize the data from treated cultures. The experiment was repeated three times in duplicates. Values plotted are mean +/- standard error. Data were analyzed by Mann-Whitney U-test. *$p<0.05$.

**SUPPLEMENTARY FIGURE-8**

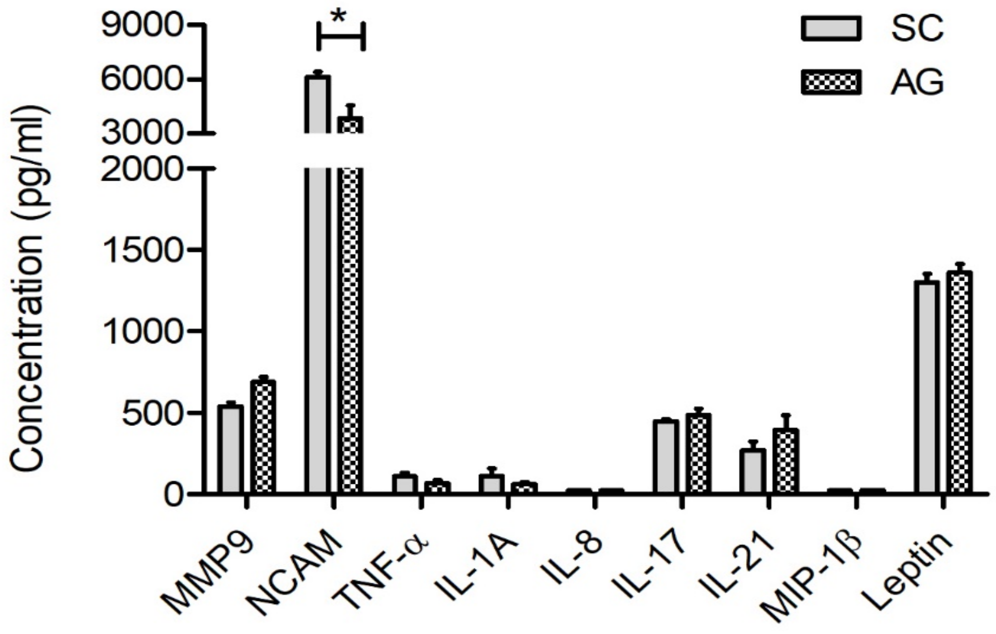


**Supplementary Figure 8.** Levels of inflammatory cytokines/chemokines in rabbit lungs infected with Mtb-SC or AG at 4 weeks. Filtered lung homogenates were used to measure the markers by ELISA. n=3 animals Per group. Values plotted are mean +/- standard error. Data was analyzed by Mann-Whitney U-test. *p<0.05.

Cellular Necrosis network gene expression

| Genes | Prediction (based on direction) | AG-vs-SC |
|---|---|---|
| ADM | Increased | -5.5 |
| NR2C2 | Increased | -4.8 |
| WEE1 | Increased | -4.6 |
| GDA | Increased | -4.3 |
| BLNK | Increased | -4.1 |
| TBX21 | Increased | -4.1 |
| CD200 | Increased | -4.0 |
| IGF1 | Increased | -3.9 |
| LAPTM4B | Increased | -3.9 |
| TP63 | Increased | -3.8 |
| NUP107 | Increased | -3.8 |
| EEF2K | Increased | -3.7 |
| NOA1 | Increased | -3.7 |
| GBA | Increased | -3.5 |
| CDC7 | Increased | -3.1 |
| NDEL1 | Increased | -3.1 |
| FSTL1 | Increased | -3.0 |
| EZH2 | Increased | -2.9 |
| PIGR | Increased | -2.7 |
| FER | Increased | -2.7 |
| CLU | Increased | -2.6 |
| PLD1 | Increased | -2.6 |
| IQUB | Increased | -2.4 |
| NCEH1 | Increased | -2.4 |
| CD44 | Increased | -2.3 |
| ITCH | Increased | -2.3 |
| SYNE1 | Increased | -2.2 |
| PSIP1 | Increased | -2.2 |
| NUP155 | Increased | -2.2 |
| ZEB1 | Increased | -2.0 |
| CDC73 | Increased | -1.9 |
| PHIP | Increased | -1.9 |
| TOP2B | Increased | -1.9 |
| SLU7 | Increased | -1.6 |
| RB1CC1 | Increased | -1.6 |
| NEK1 | Increased | -1.5 |
| ND4 | Increased | -1.5 |
| S100A8 | Increased | 1.7 |
| GNB2 | Increased | 1.8 |
| S100A9 | Increased | 1.9 |
| MT3 | Increased | 1.9 |
| LAT | Increased | 3.2 |
| KLRK1 | Decreased | -5.7 |
| SULF1 | Decreased | -4.6 |
| ATG14 | Decreased | -4.5 |
| GPR37 | Decreased | -4.5 |
| MBD4 | Decreased | -4.4 |
| ADCY10 | Decreased | -4.1 |
| NCOA2 | Decreased | -4.0 |
| BRCA1 | Decreased | -3.7 |
| WDR48 | Decreased | -3.6 |
| TFB1M | Decreased | -3.6 |
| RASSF6 | Decreased | -3.5 |

Apoptosi

| Genes |
|---|
| GPR37 |
| BRCA1 |
| CLU |
| NCOA2 |
| CADM1 |
| IGF1 |
| CREB1 |
| JMJD1C |
| UBE2B |

| | | |
|---|---|---|
| PDGFD | Decreased | -2.9 |
| SGPL1 | Decreased | -2.3 |
| SMAD4 | Decreased | -2.1 |
| SENP2 | Decreased | -2.1 |
| MED1 | Decreased | -2.0 |
| MIB1 | Decreased | -2.0 |
| ADAMTS1 | Decreased | -1.9 |
| FKBPL | Decreased | -1.8 |
| SOX4 | Decreased | 1.6 |
| CA3 | Decreased | 1.9 |
| CREB1 | Decreased | 1.9 |
| ALKBH3 | Decreased | 2.0 |
| TRAF3IP2 | Decreased | 2.8 |
| MAP3K2 | Decreased | 3.5 |
| EGLN1 | Decreased | 3.5 |
| LANCL1 | Decreased | 3.6 |
| RAB32 | Decreased | 3.9 |
| SERPINB10 | Decreased | 4.0 |
| MARVELD3 | Decreased | 4.3 |

**s network gene expression**

| Prediction (based on direction) | AG-vs-SC |
|---|---|
| Increased | -4.455 |
| Increased | -3.717 |
| Increased | -2.624 |
| Increased | -4.046 |
| Increased | -1.624 |
| Increased | -3.947 |
| Affected | 1.911 |
| Affected | -1.946 |
| Affected | 1.006 |

***Autophagy network gene expression***

| Genes | Prediction (based on direction) | AG-vs-SC |
|---|---|---|
| BRCA1 | Increased | -3.717 |
| EIF4G1 | Increased | -1.931 |
| SLU7 | Increased | -1.593 |
| GPR37 | Decreased | -4.455 |
| EEF2K | Decreased | -3.689 |
| GOPC | Decreased | -1.262 |
| PINK1 | Decreased | 1.149 |
| TP63 | Decreased | -3.832 |
| SMAD4 | Decreased | -2.146 |
| MID2 | Decreased | -2.946 |
| RB1CC1 | Decreased | -1.582 |
| NUPR1 | Decreased | 2.711 |
| ROCK1 | Decreased | -1.057 |
| PLD1 | Decreased | -2.567 |
| IGF1 | Decreased | -3.947 |
| LAPTM4B | Decreased | -3.928 |
| ATG14 | Decreased | -4.503 |
